# A novel heuristic of rigid docking scores positively correlates with full-length nuclear receptor LRH-1 regulation

**DOI:** 10.1101/2024.05.05.592617

**Authors:** Zeinab Haratipour, David Foutch, Raymond D. Blind

## Abstract

The nuclear receptor Liver Receptor Homolog-1 (LRH-1, *NR5A2*) is a ligand-regulated transcription factor and validated drug target for several human diseases. LRH-1 activation is regulated by small molecule ligands, which bind to the ligand binding domain (LBD) within the full-length LRH-1. We recently identified 57 compounds that bind LRH-1, and unexpectedly found these compounds regulated either the isolated LBD, or the full-length LRH-1 in cells, with little overlap. Here, we correlated compound binding energy from a single rigid-body scoring function with full-length LRH-1 activity in cells. Although docking scores of the 57 hit compounds did not correlate with LRH-1 regulation in wet lab assays, a subset of the compounds had large differences in binding energy docked to the isolated LBD *vs*. full-length LRH-1, which we used to empirically derive a new metric of the docking scores we call “ΔΔG”. Initial regressions, correlations and contingency analyses all suggest compounds with high ΔΔG values more frequently regulated LRH-1 in wet lab assays. We then docked all 57 compounds to 18 crystal structures of LRH-1 to obtain averaged ΔΔG values, which robustly and reproducibly associated with full-length LRH-1 activity in cells. Network analyses on the 18 crystal structures of LRH-1 suggest unique communication paths exist between the subsets of LRH-1 crystal structures that produced high *vs.* low ΔΔG values, identifying a structural relationship between ΔΔG and the position of Helix 6, a previously established regulatory helix important for LRH-1 regulation. Together, these data suggest computational docking can be used to quickly calculate ΔΔG, which positively correlated with the ability of these 57 hit compounds to regulate full-length LRH-1 in cell-based assays. We propose ΔΔG as a novel computational tool that can be applied to LRH-1 drug screens to prioritize compounds for secondary screening.

**Graphical Abstract:**
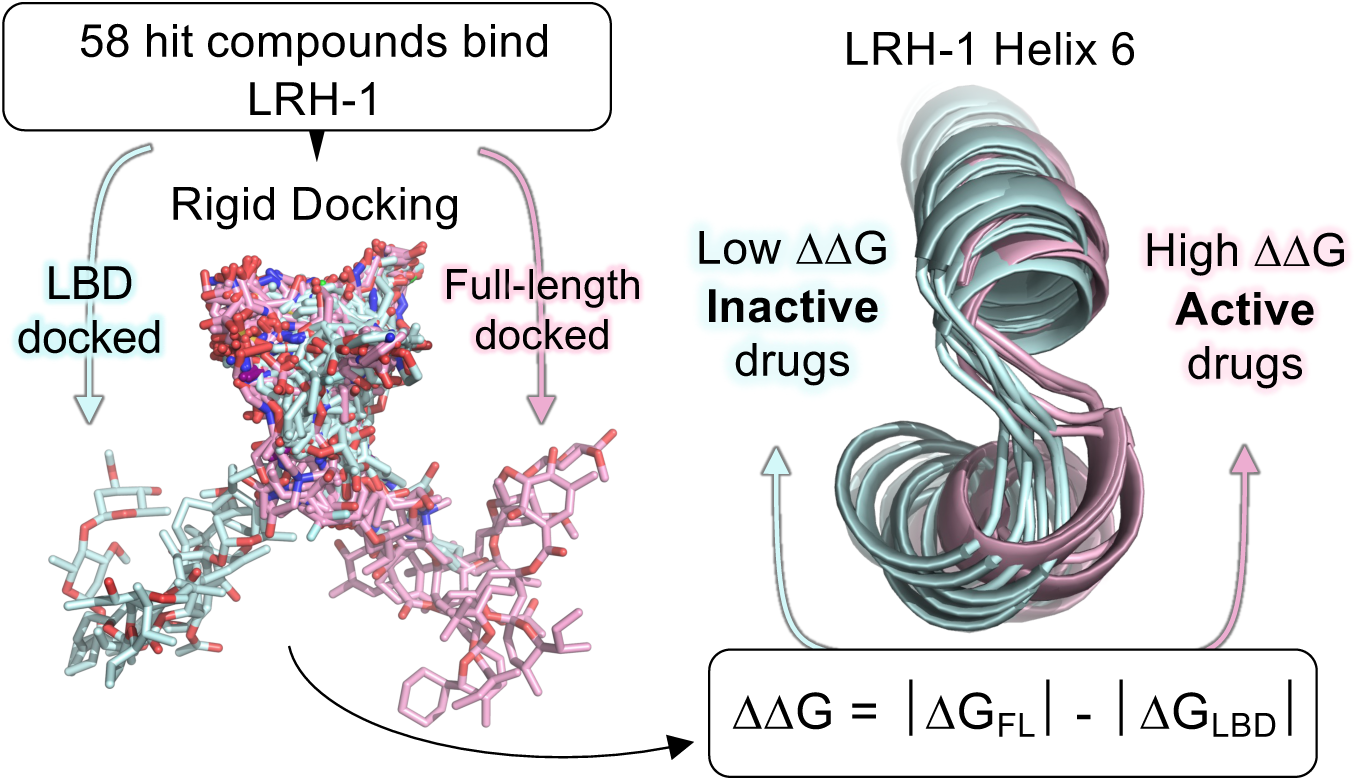

## Introduction

Prioritizing hit compounds identified from a primary compound screen is important for drug development[1–6] as follow-up secondary assays are resource-intensive, particularly cell-based assays of target protein function[7,8]. Cell-based assays monitor compound activity in a physiologically relevant environment, increasing confidence in results[7,9–11] while retaining compatibility with higher-throughput screening formats[12]. The choice of secondary assay must therefore strike a balance between testing fewer, high-priority compounds in a higher-confidence assay, or more compounds in a lower-confidence assay, since resources not unlimited[13]. Computational docking has been used to help prioritize compounds[14], which can improve the hit rate from secondary assays[2–6,15]. However, little wet-lab data are available validating direct relationships between docking scores and functional regulation in cells[16–18], particularly for non-enzyme allosterically-regulated target proteins[19,20]. Still, the speed, accessibility and low cost of computational docking provides an attractive way to prioritize compounds[21,22]. Thus, there is a need for docking approaches that have been validated with wet-lab data, correlating docking scores with compound activity in cell-based assays[2–4,23,24] particularly for allosterically-regulated targets[5,20,22]. Here, we discovered and validated one such approach, retrospectively correlating published cell-based functional data[1] with new rigid-body docking of compounds to Liver Receptor Homolog-1 (LRH-1, *NR5A2*), an allosterically regulated nuclear receptor and drug target for several human diseases.

The nuclear receptor superfamily is a group of DNA-binding transcription factors regulated by hydrophobic ligands such as cholesterol-based steroids[25], fatty acids[26], heme-based metabolites[27] and phospholipids[28,29]. LRH-1 (*NR5A2*) is a nuclear receptor expressed ubiquitously in humans[30,31], but is particularly important for adult function in the liver[32–37], pancreas[38,39] and gut[40–42]. A wide variety of physiological studies have provided pre-clinical validation of LRH-1 as a drug target in diabetes and non-alcoholic fatty liver disease (NAFLD)[33,36], pancreatic cancer[43–45] and ulcerative colitis[42], all diseases with unmet clinical need. Although there has been great recent progress in the development of LRH-1 regulatory compounds[42,46–49], no FDA-approved drugs currently target LRH-1, so there remains need for novel approaches that can hasten LRH-1 drug development.

Nuclear receptor ligands bind to a canonical cleft in the ligand binding domain (LBD), which alters the conformation of the LBD, which permits interaction with a variety of transcriptional co-regulator proteins[50–52]. This principle applies to LRH-1 and the highly homologous SF-1 nuclear receptors, which are both regulated through this canonical mechanism by several ligands, including natural phospholipids[37,42,53–62] and synthetic small molecules[37,42,46–48,63–65]. These regulatory ligands all bind LRH-1 with 1:1 stoichiometry, as shown in many different crystal structures of human LRH-1 in the protein data bank (PDB). The allosteric mechanism of ligand-induced interaction with coregulator proteins is utilized in LRH-1 drug screening platforms such as AlphaScreen[66], in which compounds induce a measured interaction between LRH-1 and a small peptide that represents a transcriptional coregulator protein [67–69], such as TIF2[70] or PGC1α[71]. These screens, however, can only identify compounds that regulate LRH-1 through the canonical coregulator-recruitment mechanism, and are not able to identify any compounds that bind and regulate LRH-1 through non-canonical mechanisms.

To address this, we executed an LRH-1 compound screen that identified 58 new compounds which simply bind directly to LRH-1 from the 2322 compound Spectrum Discovery library[1]. This FRET-based screen measured the ability of each compound to compete with a probe installed in the canonical ligand binding site of LRH-1, suggesting these compounds also bind LRH-1 at the canonical ligand binding site. The hit compounds were subjected to several secondary assays monitoring functional regulation of LRH-1, which showed 14 of the 58 compounds regulated either 1) the isolated LBD in a coregulator binding assay or 2) full-length LRH-1 in a luciferase reporter assay driven by the *CYP8B1* promoter or 3) the *CYP17A1* promoter, both established LRH-1 target promoters in cells. Surprisingly and against the standard dogma of nuclear receptor regulation, there was almost no overlap in these compounds. The compounds either regulated coregulator binding to the LBD, or the compounds regulated full-length LRH-1 assays in cells, with no overlap, even though these compounds were all identified based on their ability to directly bind pure, recombinant LRH-1[1].

Here, we examined the interaction of these compounds with LRH-1 by applying PyRx computational rigid docking[72] to predict relative docked binding energies to a total of 19 structural models of LRH-1 (18 crystal structures of the isolated LBD and 1 integrative model of the full-length LRH-1[73]). We then correlated those docking results retrospectively with published cell-based functional data from our previous study (439 total functional assay measurements)[1]. We found that docking to the full-length model of LRH-1[73] predicts different binding positions for compounds active on full-length LRH-1, which were closer to the entrance of the ligand binding pocket and Helix 6, an important helix that helps mediate ligand activation of LRH-1 [37,64,70,71,74–76]. The scores of compounds docked to full-length LRH-1 or the isolated LBD did not correlate with LRH-1 regulation in wet lab assays, however several compounds had large differences in docking scores (ΔG) to the full-length *vs*. isolated LBD of LRH-1, an arbitrary metric we call ΔΔG. We found ΔΔG positively associated with the ability of our set of 58 hit compounds to regulate full-length LRH-1 in cell-based assays. Network analyses suggest the position of Helix 6 as a structural element that may contribute to the relationship between ΔΔG and ligand-mediated LRH-1 regulation in cells, consistent with analyses by independent groups[37,64,70,71,74–76]. The data presented here suggest a role for Helix 6 in ligand-mediated regulation of LRH-1, similar to other studies which have applied comparative crystallography, MD simulations and network analyses[37,64,70,71,74–76]. The ΔΔG metric can help prioritize hit compounds for expensive secondary screening, which can hasten LRH-1 compound development. ΔΔG can be rapidly generated even for large hit lists, as minimal resources are required for rigid-body PyRx-based docking, highlighting practical utility.

## Results

### Compounds which functionally regulated the isolated LBD, did not regulate full-length LRH-1 in cells

We previously identified 58 compounds that bind directly to the ligand binding domain (LBD) of LRH-1 (**Fig 1A**) using a FRET-based competition screen[1] (**Fig 1B**). We then applied three independent high-throughput assays to test these 58 compounds for functional regulation of either the purified recombinant LBD in a coregulator binding screen (98 independent assay measurements) or full-length LRH-1 regulation in cells (341 independent assay measurements), for a total 439 independently measured events of compound-induced regulation of LRH-1 function[1] (**Supplemental Spreadsheet 1**). Regulation of the LBD was determined by fluorescence anisotropy (**Fig 1C**), which measured compound-induced interaction between the isolated LBD of LRH-1 and a peptide representing a known transcriptional coregulator of LRH-1[71]. Full-length LRH-1 regulation in cells was determined by measuring compound-induced LRH-1 activation of a luciferase reporter gene, driven by one of two well-established LRH-1 promoter DNA sequences (*CYP8B1* or *CYP17A1*) in HEK293T cells (**Fig 1D**). We showed of the 58 compounds identified to bind LRH-1, 14 compounds regulated LRH-1 function in at least one of these three assays[1] (Dunnett’s corrected *p_adj_*<0.05, **Supplemental Spreadsheet 2**). Here, we applied principal component analysis (**Fig 1E**) of the equally scaled log_2_ fold-change data from all 439 wet lab assay measurements, suggesting distinct patterns of regulation for the isolated LBD *vs*. full-length LRH-1 assays, despite that all these compounds directly bind LRH-1[1]. Contingency analyses suggest compounds that significantly regulated the isolated LBD, less frequently regulated full-length LRH-1 in cells (**Fig 1E**, 90% vs. 7%, *p*<0.0001 by Fisher’s exact), and compounds that regulated full-length LRH-1 in cells less frequently regulated the isolated LBD (**Fig 1F**, 68% vs. 8%, *p*=0.0005 by Fisher’s exact). Together these data suggest compounds that regulated the isolated LBD did not regulate full-length LRH-1 in cells, despite that all the compounds were identified based on their ability to directly bind LRH-1 (**Fig 1B**). We next used computational docking to determine if there were any differences in the docking scores (docked binding energies) of these compounds to the isolated LBD *vs*. full-length models of LRH-1.

**Figure 1.**
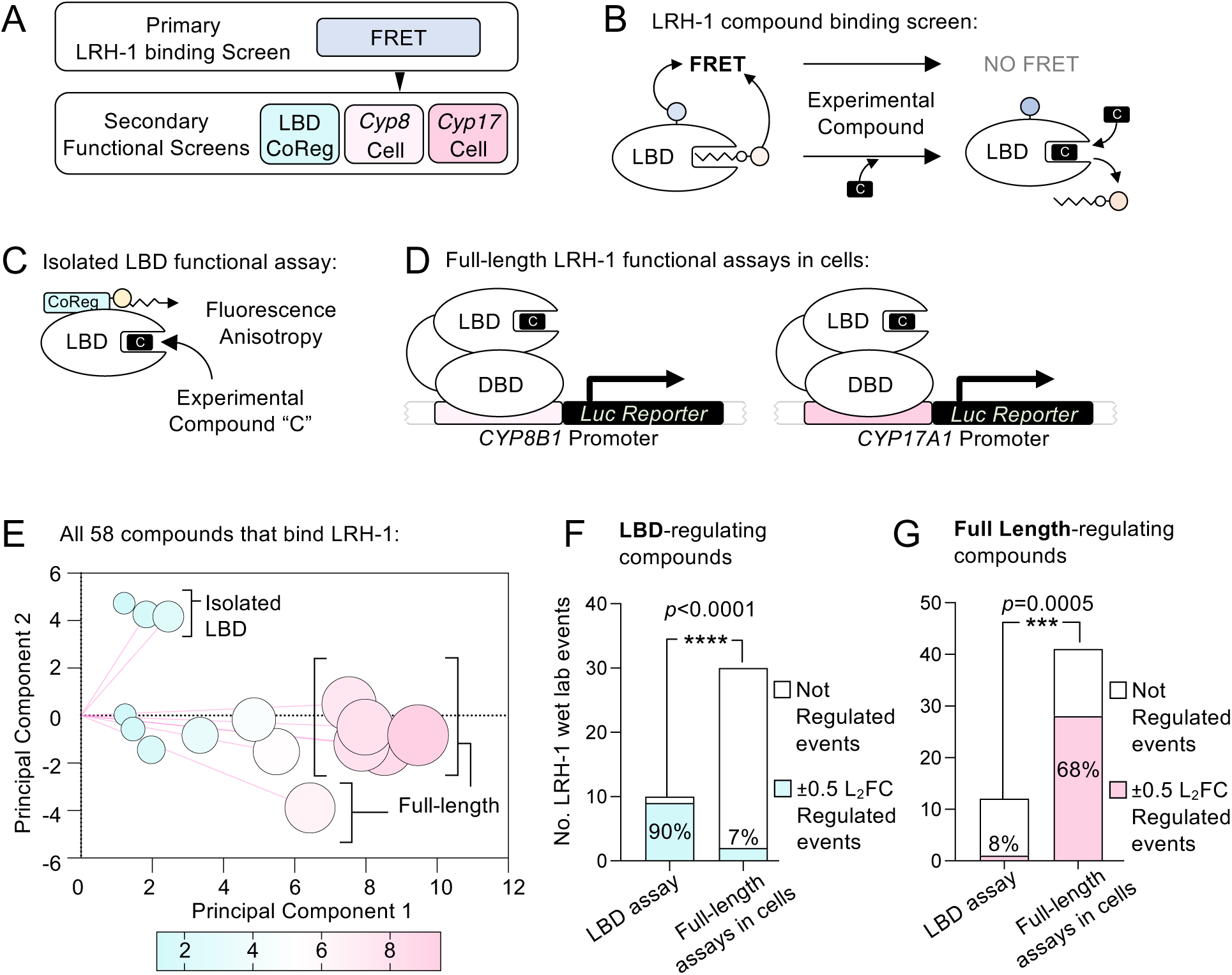
Compounds identified to bind LRH-1 LBD from a previous wet-lab screen regulated either the ligand-binding domain (LBD) or full-length LRH-1 in cells, with little overlap. **A.** Previously published screening strategy highlighting **B**. primary screen that identified 58 compounds based on their ability to compete with a FRET probe installed at the canonical ligand binding site of the isolated ligand binding domain (LBD) of human LRH-1. **C.** Schematic of isolated LBD assay that measured compound-induced interaction between PGC1α coregulator peptide and the isolated LBD of LRH-1. **D**. Schematic of full-length LRH-1 luciferase reporters in HEK293T cells using either the *CYP17A1* or *CYP8B1* promoters. **E**. Biplot of principal component analysis of all 439 wet-lab assayed events (assays from panels C-D) induced by all 58 LRH-1 hit compounds, suggesting the isolated LBD assay *vs*. full-length LRH-1 assays do not cluster together. **F**. Contingency analysis of the frequency of LRH-1 regulation induced by all compounds that regulated the isolated LBD assay (5 compounds, 40 assays), showing LBD-regulating compounds less frequently regulated full-length LRH-1 in cells (90% vs. 7%, *p*<0.0001 by Fisher’s exact test), regulation defined as *p_adj_*<0.05 Dunnett’s corrected and log_2_ fold change at least ±0.5. **G**. Contingency of LRH-1 regulation by the compounds that regulated full-length LRH-1 in cells (9 compounds, 53 assays), showing full-length regulating compounds less frequently regulated the isolated LBD assay (68% vs. 8%, *p*=0.0005 by Fisher’s exact). *These data suggest that even though all these compounds were identified based on their ability to bind the isolated LBD, compounds that regulated the isolated LBD did not regulate the full-length LRH-1 in cells as frequently, and vice versa*.

### Rigid-body compound docking to full-length LRH-1 vs. the isolated LBD yields divergent binding positions and binding energies

Our previous study docked all 2322 compounds to a crystal structure of the isolated ligand binding domain (LBD) of LRH-1 (PDB:6OQX), which showed no correlation between the docking scores and regulation of LRH-1[1]. Here, we used the same PyRx rigid-body docking approach, but now examined docking to the full-length LRH-1 model (PDB_DEV: 00000035, **Fig 2A, Supplemental Spreadsheet 3**). Importantly, the model of full-length LRH-1 is a solution-based, integrated structure that was extensively computationally optimized by Rosetta and several other approaches[73]. We therefore expected ligand docking to this model to result in lower binding energies when compared to a crystallographically restrained model of the isolated LBD. Indeed, we observed significantly lower docked binding energies for compounds docked to full-length LRH-1 *vs*. the isolated LBD (**Fig 2B**). The docked positions of the compounds were also different (**Fig 2C**), specifically the 5 compounds that regulated the isolated LBD (**Fig 1C**) docked within the well-established, canonical ligand binding site, deep in the core of the LRH-1 protein (**Fig 2D**). However, some of the 9 compounds that regulated full-length LRH-1 in cells (**Fig 1D**) docked to the full-length LRH-1 at the entrance to the canonical ligand biding site, clustered around Helix 6 (**Fig 2E**). Our previous work suggests all 9 of these compounds (**Fig 2E**) bind directly to the isolated LRH-1 LBD, as well as regulate full-length LRH-1 function in cells[1]. Further, we determined saturable binding constants for two compounds that docked at the entrance of the ligand binding site near Helix 6, VU0243218 (IC_50_=9.4μM, 95%CI 8.1-10.8μM) and VU0656021 (IC_50_=27.0μM, 95%CI 16.1-171.5μM), again suggesting direct interactions of these compounds with the LRH-1 ligand binding domain (**Fig 2E**). Full transcriptomes induced by 10μM VU0243218 showed selective regulation of endogenous ChIP-seq target genes of LRH-1, and a specific chemical competitor of LRH-1 attenuated the gene expression response of endogenous LRH-1 target genes to VU0243218[1]. This level of validation could not be applied to all hit compounds from the screen, however the data suggest these compounds indeed bind and regulate LRH-1, despite docking at the entrance to the ligand binding pocket. No crystal structures have been solved for any of these recently identified LRH-1 compounds, thus the precise binding mode for the compounds awaits those studies. These data suggest ligands that bind and regulate the isolated LBD *vs*. full-length LRH-1 in cells have different rigid-body docking to the isolated LBD *vs*. full-length models of human LRH-1. Since these are the first computational ligand docking studies to the full-length LRH-1[73], we next sought to determine if docked binding energy to full-length LRH-1 correlated with the ability of compounds to regulate LRH-1.

**Figure 2.**
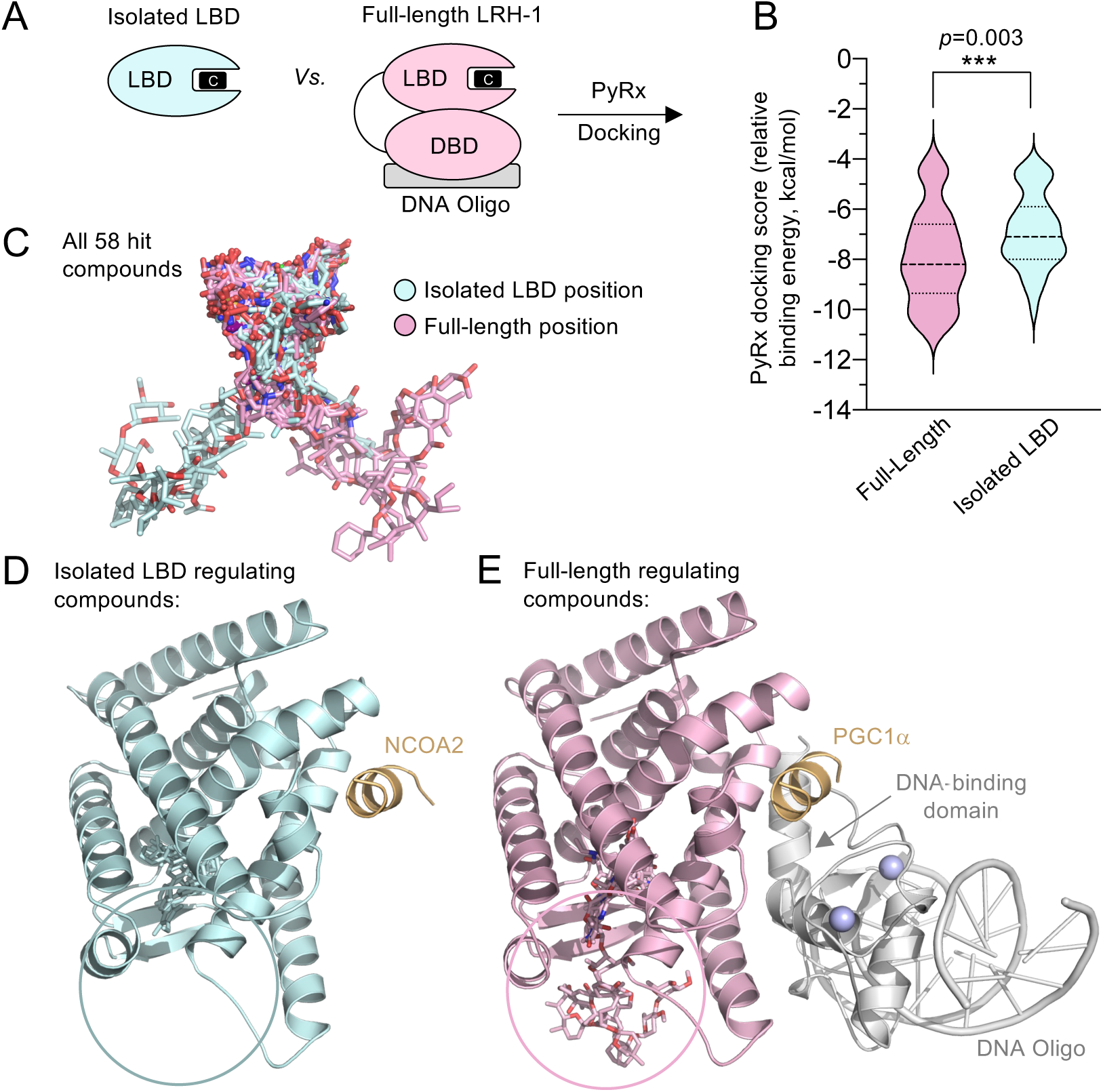
The 58 hit compounds computationally docked to full-length LRH-1 and the isolated LBD differently. **A**. Schematic showing differences in isolated LBD vs. full length LRH-1 used for PyRx rigid docking. **B.** Plot of docking scores (relative binding energies, kcal/mol) of the 58 hit compounds docked to the isolated LBD (PDB:6OQX, teal) *vs*. full-length LRH-1 (PDB_DEV: 00000035, pink), showing lower binding energy of compounds to full-length LRH-1, \*\*\**p*=0.0003 by two-tailed paired t-test. **C**. Close up of the different docked positions of all 58 hit compounds docked to the isolated LBD (PDB:6OQX, teal) vs. full-length LRH-1 (pink), with the protein removed for clarity. **D.** Docked positions in LRH-1 LBD (PDB:6OQX) of only the 5 hit compounds that regulated the isolated LBD, co-crystalized NCOA2 peptide (gold) is shown for orientation. **E**. Docked positions of only the 9 hit compounds that regulated the full-length LRH-1 in cells, DNA oligo shown in gray, Zinc atoms purple spheres, PGC1α peptide shown in gold. *These data suggest compounds dock to the isolated LBD vs. full-length LRH-1 models slightly differently*.

### Compound ΔΔG associates with the ability of compounds to regulate full-length LRH-1 in cells

We previously established that the relative docking scores of compounds to the isolated ligand binding domain of LRH-1 (PDB:6OQX) did not correlate with regulation of full-length LRH-1 in cells[1]. However, given the different docking scores we observed (**Fig 2B**), we asked if docked binding energy to full-length LRH-1 associated with the ability of compounds to regulate full-length LRH-1 in cells. Since the hit compounds both activated and inhibited LRH-1 in the wet lab assays, the data contain both negative (inhibitory) and positive (activating) log2 fold change values for each compound, compared to DMSO controls (**Supplemental Spreadsheet 1**). Thus, we converted these continuous variables to discrete variables by applying log2 fold change cutoffs to define discrete “regulated events” *vs*. “not regulated events” and treated each wet lab measurement of LRH-1 function (each well in the high throughput assays) as an independent event. We analyzed the continuous data (**Fig 6**) and present these discrete analyses first. Plotting the binding energy of each compound (*Δ*G) docked to either the isolated LBD of LRH-1 (PDB:6OQX, **Fig 3A**) or full-length LRH-1 (**Fig 3B**) as a function of the discrete number of wet-lab events regulated by each compound, and using a very inclusive cutoff for regulation (±0.25 log_2_fold change *vs.* DMSO control) failed to show a correlation by Spearman rank, or a non-zero slope of a linear regression by F-test, for either the isolated LBD (*p*=0.48) or full-length LRH-1 (*p*=0.52). These data suggest the relative docked binding energy of compounds to full-length LRH-1 did not correlate with the ability of that compound to regulate LRH-1 in the assays examined here. However, we noted some compounds had large differences in binding energy to the isolated LBD vs. full-length LRH-1, and those compounds appeared to more frequently regulate full-length LRH-1 in the wet lab assays. Thus, we empirically derived the heuristic ΔΔG metric, which is simply the absolute value of the PyRx docking score (relative docked binding energy for each compound) to the full-length LRH-1 (*Δ*G_FL_), less the absolute value of the *Δ*G for each compound binding to the isolated LBD of LRH-1 (*Δ*G_LBD_, **Fig 3C**). We plotted *ΔΔ*G against the identical wet-lab functional data, which showed a slight upward trend to the regression when the 6OQX crystal structure was used to calculate *ΔΔ*G_6OQX_, however the slope was not significantly non-zero (**Fig 3D**). We tested this further by docking to a different crystal structure of the LBD (PDB:1YOK) to derive the *ΔΔ*G metric, which produced a significant Spearman rank correlation (r=0.291, \**p*=0.014) and a slightly non-zero slope to a linear regression (\**p*=0.043 by F-test). We excluded one compound from these analyses (VU0656093), as it was the only compound to produce positive (unfavorable) binding energies docked to PDB:1YOK (+15.3kcal/mol), full-length LRH-1 (+26.6kcal/mol), and PDB:6OQX (+44.1 kcal/mol), more that 11-fold higher than the next highest binding energy compound (**Supplemental Spreadsheet 3**). Thus this compound was excluded and only 57 compounds were analyzed for the remainder of this study. These initial data suggested a weak, but perhaps non-zero association might exist between *ΔΔ*G_1YOK_ full-length LRH-1 regulation in cell-based assays. Since any computational approach that could improve cell-based secondary screens would have great practical value in larger LRH-1 drug screens in both large pharmaceutical companies and within academia, we followed up on this initial observation.

**Figure 3.**
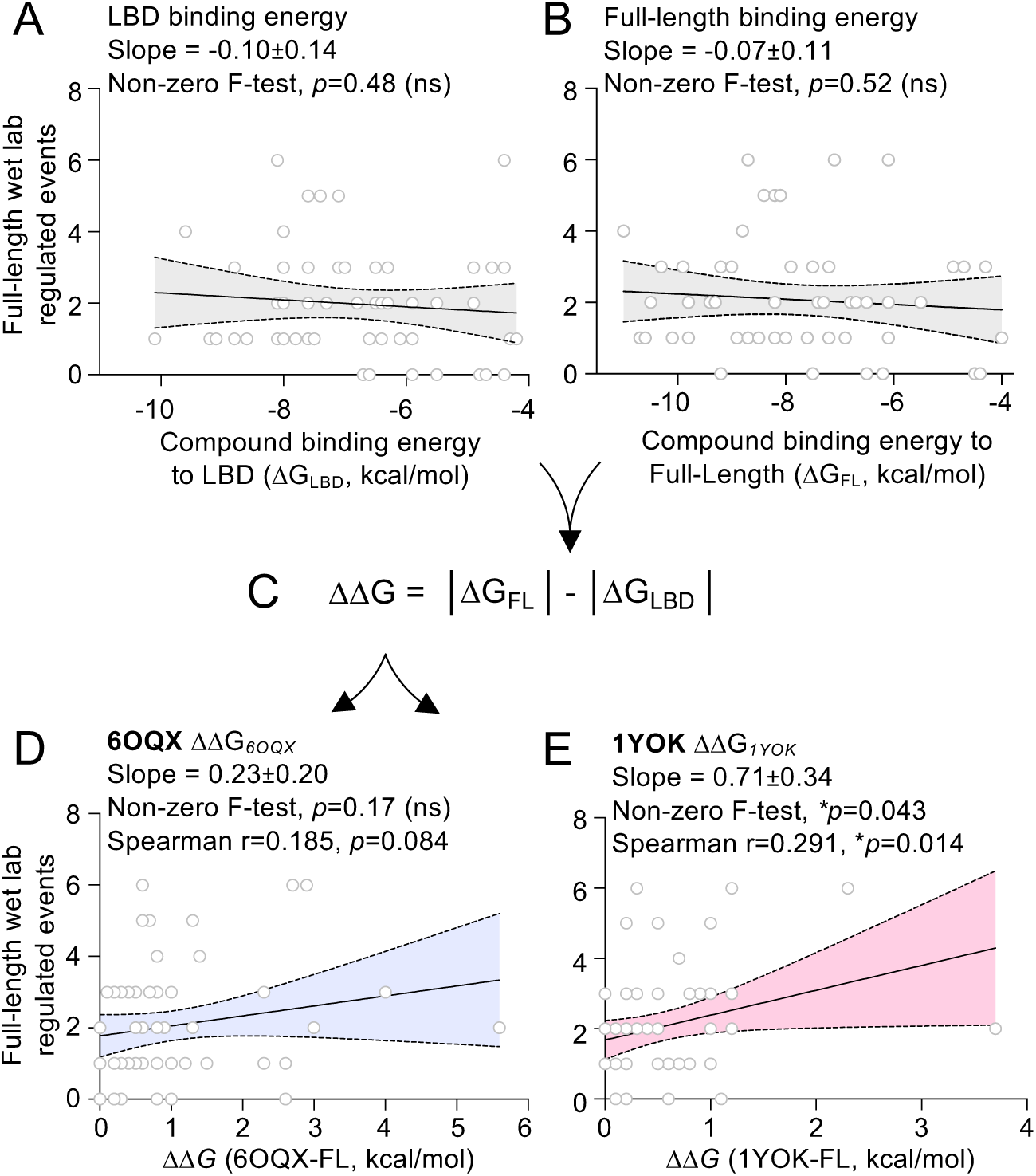
Relative binding energy (ΔG) of compounds docked to either the ligand binding domain or full-length LRH-1 did not associate with LRH-1 regulation in cells. **A**. Plot of docked binding energy of 57 compounds to the isolated ligand binding domain (LBD, PDB:6OQX) *vs*. the number of wet lab assay events regulated by each compound (full-length LRH-1 regulation of the *CYP17A1* and *CYP7B2* luciferase promoters), with each of the wells in the high-throughput assays treated as an individual event (replicates not averaged) and regulation defined as meeting ±0.25 log_2_ fold change cutoff *vs*. DMSO control. Solid line indicates linear regression for all points, shaded area is 95% confidence of regression, slope of the regression and F-test *p* value for non-zero slope indicated, with no correlation by Spearman rank correlation. **B**. Identical as A, but plotting docked binding energy of compounds to full-length LRH-1 (PDB_DEV: 00000035). **C**. ΔΔG is an arbitrary metric based on the difference in docked binding energy of a compound to the full-length LRH-1 (ΔG_FL_) *vs*. the isolated LBD (ΔG_LBD_). **D**. Plot of *ΔΔ*G calculated using PDB:6OQX (*ΔΔ*G*_6OQX_*) *vs*. the number of wet lab assay events regulated by each compound (full-length LRH-1 regulated events from the *CYP17A1* and *CYP8B1* luciferase assays). **E**. Identical as D, but ΔΔG calculated using docking scores to LRH-1 structure 1YOK (ΔΔG*_1YOK_*), showing slightly non-zero slope and significant one-tailed Spearman correlation, calculated in Prism. *These data suggest ΔΔG values of compounds could correlate with wet lab activity on LRH-1*.

### Averaged ΔΔG across 18 crystal structures of LRH-1 associates with the ability of a compound to regulate LRH-1 in all wet-lab assays

We docked the 57 hit compounds to 18 crystal structures of the human LRH-1 ligand binding domain (see methods for the list of all 18 structures from the protein data bank, PDB) and calculated *ΔΔ*G values for all 57 compounds to each of the 18 structures of LRH-1, which generated a matrix of 1026 *ΔΔ*G values (**Supplemental Sheet 4**). We averaged the 18 *ΔΔ*G values for each compound (one *ΔΔ*G value for each of the 18 crystal structures) and plotted this “averaged *ΔΔ*G” (referred to as *ΔΔ*G for the remainder of this manuscript) for each compound as a function of the number of wet-lab events regulated by that compound. Spearman correlations were statistically significant, and slopes of linear regressions were non-zero at ±0.25 L_2_FC (**Fig 4A**), ±0.5 L_2_FC (**Fig 4B**) or ±1.0 L_2_FC (**Fig 4C**) cutoffs, again suggesting an association between *ΔΔ*G and the ability of a compound to regulate LRH-1 function in wet-lab assays. Contingency analyses further suggested compounds with higher *ΔΔ*G values were significantly more likely to functionally regulate LRH-1 (**Fig 5A-F**). Specifically, compounds with a *ΔΔ*G value higher than 1.0 kcal/mol regulated LRH-1 in 23% of all wet lab measured events, whereas compounds with *ΔΔ*G below 1.0 kcal/mol regulated LRH-1 in only 9% of all wet lab events (**Fig 5A**, *p*=0.0001 by Fisher’s exact test). Compounds in the highest 10^th^ percentile of *ΔΔ*G values regulated LRH-1 in 30% of wet lab events, while all remaining compounds regulated LRH-1 in only 11% of wet lab events (**Fig 5B**, *p*=0.0014). Compounds in the highest quartile of *ΔΔ*G values regulated LRH-1 in 24% of wet lab events, all remaining compounds regulated LRH-1 in only 9% of wet lab events (**Fig 5C**, *p*=0.0002). The highest 3 quartiles of *ΔΔ*G values also selectively regulated LRH-1 in 16% of wet lab events, while all remaining compounds regulated LRH-1 in only 4% of wet lab events (**Fig 5B**, *p*=0.0014). This association was significant if a more rigorous cutoff of ±1.0 log_2_ fold change was used (**Fig 5E-F**). Together these data suggest that among these 57 hit compounds, compounds with higher values of *ΔΔ*G more frequently associated with regulation of LRH-1 in the wet lab.

**Figure 4.**
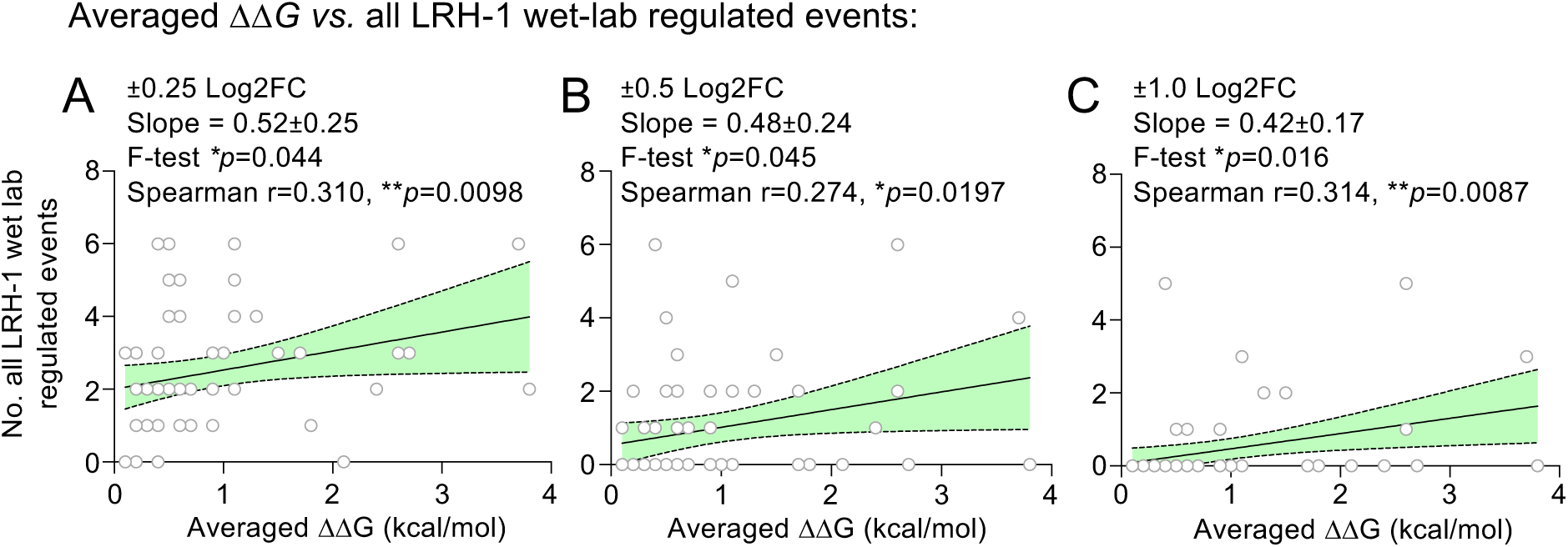
Linear regression of ΔΔG averaged across all 18 crystal structures of LRH-1 plotted vs. the number of compound-regulated LRH-1 assay events has a non-zero slope. **A**. 57 of 58 hit compounds (one compound excluded due to high positive binding energies docked to several LRH-1 models, see methods) were docked to 18 crystal structures of the human LRH-1 LBD and ΔΔG values calculated, producing a matrix of 1102 ΔΔG values, provided in supplemental data. The average of the 18 ΔΔG values (one value for each of the 18 crystal structures) for each of the 57 compounds were plotted as a function of the number of all LRH-1 wet-lab regulated events induced by that compound, with regulation discretely defined as at least ±0.25 Log2 fold change from DMSO control, **B.** ±0.50 Log2FC or **C**. ±1.0 Log2FC. For all plots solid line indicates linear regression for all points, shaded area is 95% confidence interval of regression, slope and *p* value from an F-test for a non-zero value of the slope, r and p values for one-tailed Spearman correlation indicated. Spearman correlations were one-tailed as positive values of initial linear regressions suggested a positive correlation, making a negative correlation unlikely. *These data suggest compounds with high averaged ΔΔG might associate with LRH-1 activity*.

**Figure 5.**
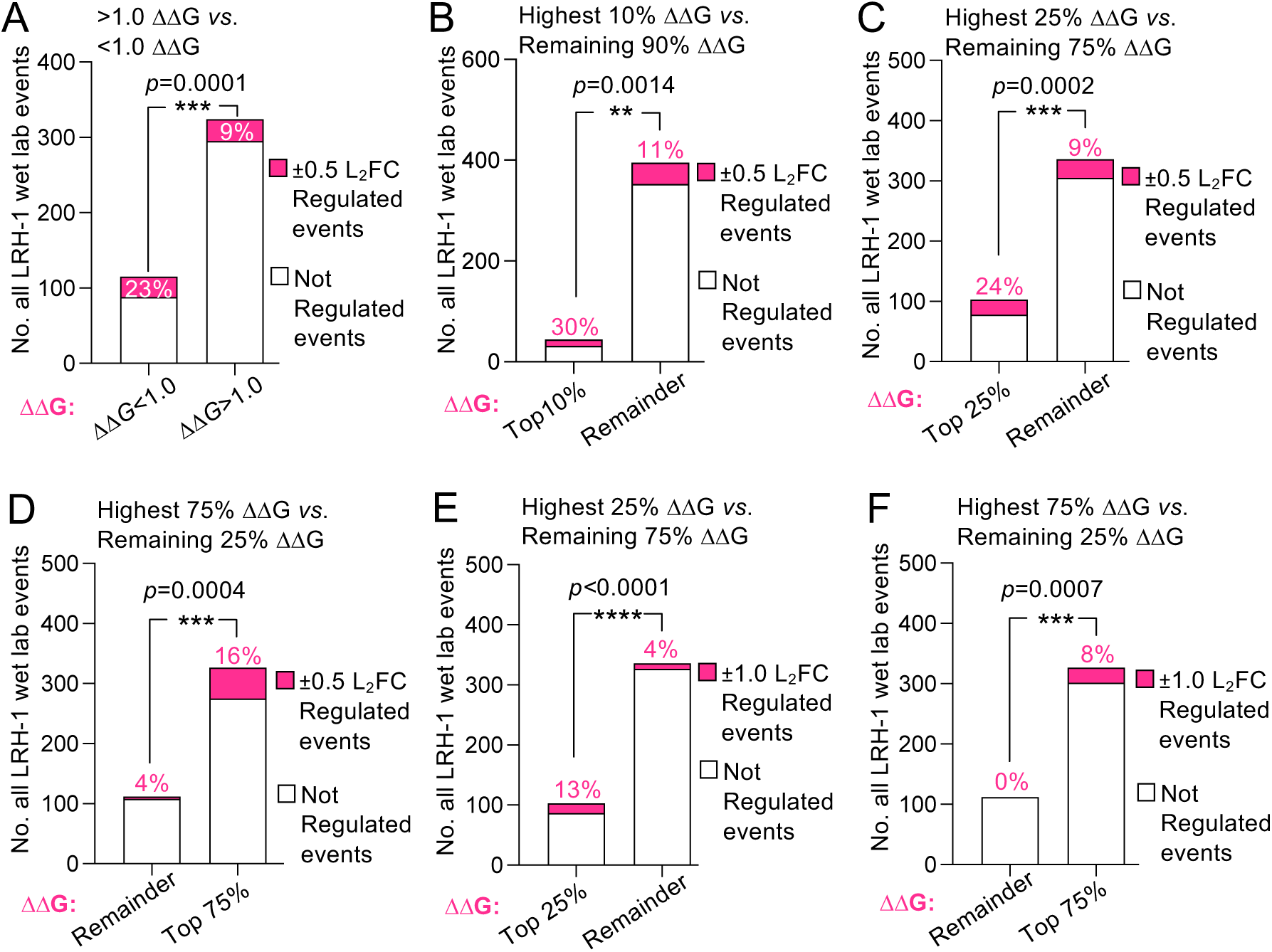
Compounds with high ΔΔG values more frequently regulated LRH-1 in wet lab assays. Contingency analyses of the frequency of LRH-1 regulation induced by all 57 compounds in all 439 wet lab assayed events (y-axis), with each well in the high-throughput assays treated as an individual event. Since compounds both inhibited and activated LRH-1, regulation in each well was defined as meeting the indicated Log_2_ fold change cutoff (L_2_FC) compared to DMSO control. **Pink section** of bars is the number of events that met the L_2_FC cutoff (percentages indicate percentage of regulated events), **white section** of bars is the number of events that did not meet L_2_FC cutoff. **A**. Contingency analysis of the frequency of LRH-1 regulation in all 439 assays, comparing compounds with ΔΔG values lower than 1.0 (*ΔΔG*<1.0) *vs*. higher than 1.0 (ΔΔG>1.0), showing compounds with higher ΔΔG values more frequently regulated LRH-1 (16% vs. 7%) by Fisher’s exact test (*p*=0.0001). **B.** Contingency comparing compounds with ΔΔG values in the top 10^th^ percentile (Top 10%) of all 57 ΔΔG values *vs*. all other compounds (Remainder), showing compounds with higher ΔΔG values more frequently regulated LRH-1 (30% vs. 11%, *p*=0.0014 by Fisher’s exact). **C**. Contingency showing compounds in the highest quartile (Top 25%) of ΔΔG values more frequently regulated LRH-1 (*p*=0.0265 by Fisher’s exact). **D**. Contingency showing compounds in the highest 3 quartiles (Top 75%) of ΔΔG values more frequently regulated LRH-1 (*p*=0.0004). **E**. Contingency showing compounds in the Top 25% of ΔΔG values more frequently regulated LRH-1 at a more exclusive L_2_FC cutoff (±1.0 L_2_FC, *p*=0.0012). **F**. Contingency showing compounds in the Top 75% of ΔΔG values more frequently regulated LRH-1 at a more exclusive ±1.0 L_2_FC cutoff (*p*=0.0007). *These data suggest ΔΔG positively associated with the ability of compounds to regulate LRH-1 in the wet lab assays we measured*.

### Higher ΔΔG compounds associated with full-length LRH-1 regulation in cells

We next asked if *ΔΔ*G for each compound would associate with regulation of full-length LRH-1 in cells. Indeed, plotting *ΔΔ*G for each compound as a function of only the full-length LRH-1 cell-based assay data showed a significantly non-zero slope to the linear regression at all cutoffs tested (**Fig S2A**). When *ΔΔ*G was plotted as a function of only the isolated LBD assays the slope of the linear regression failed to be significantly non-zero at all cutoffs tested and was not significantly correlated by Spearman (**Fig S2B**). Contingency analyses of the full-length LRH-1 assays (**Fig 6**) suggests compounds in the highest 10^th^ percentile of *ΔΔ*G values regulated LRH-1 in 25% of full-length LRH-1 cell-based assay events, all remaining compounds regulated full-length LRH-1 in only 3% of events (**Fig 6A**, *p*<0.0001). The increased frequency of full-length LRH-1 regulation in cells held if more inclusive cutoffs of ±0.5 L_2_FC (*p*=0.0001) or ±0.25 L_2_FC (*p*<0.0007) were used (**Fig 6A**). Compounds in the top quartile of *ΔΔ*G values using the more exclusive ±1.0 L_2_FC cutoff again showed higher *ΔΔ*G values associated with more frequent regulation of full-length LRH-1 in cells (**Fig S3A**, *p*=0.0002), although more inclusive L_2_FC cutoffs of ±0.5 and ±0.25 did not reach significance (**Fig S3A**). Compounds in the top 3 quartiles of *ΔΔ*G values also more frequently regulated full-length LRH-1 in cells, regardless of L_2_FC cutoff applied (**Fig S3B**). We then tested if simply the docking score (relative docked binding energy) to full-length LRH-1 (*Δ*G_FL_) would associate with a compound’s ability to regulate full-length LRH-1 in cells. Contingency analyses using the same full-length LRH-1 cell-based assay data found no association between compound binding energy to full-length LRH-1 (*Δ*G_FL_) and the ability of the 57 compounds to regulate LRH-1 in cells (**Fig 6B**), regardless of L_2_FC cutoff (**Fig S4A**) or how the compounds were grouped (**Fig S4B**). We also confirmed the 57 compounds had no preference to regulate either the isolated LBD or full-length LRH-1 assays in cells when the compounds were considered together, e.g., the 57 compounds together regulated 6% of isolated LBD assays and 6% of all full-length LRH-1 assays in cells (**Fig S5**). Together, these data suggest the group of compounds with higher *ΔΔ*G values more frequently regulated full-length LRH-1 in cells.

**Figure 6.**
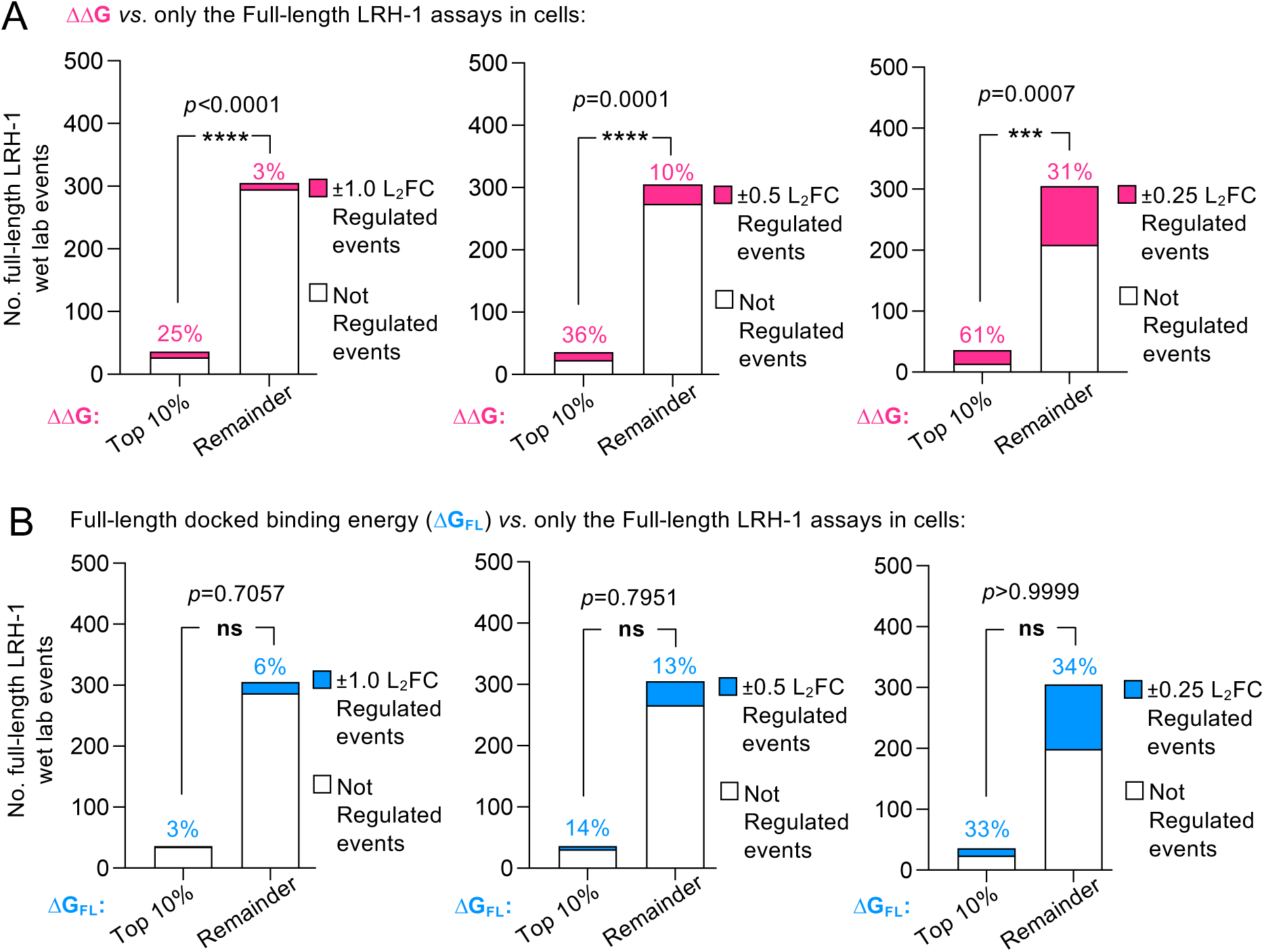
Compounds with high ΔΔG values more frequently regulate of full-length LRH-1 in cells. **A**. Contingency analysis of the frequency of full-length LRH-1 regulation in cells induced by compounds in the top 10^th^ percentile (Top 10%) of all 57 ΔΔG values *vs*. all other compounds (Remainder), examining only the 341 full-length LRH-1 assayed events in cells (isolated LBD assay results were excluded), percentages indicate the percentage of regulated (pink) events within that group of compounds, at indicated L_2_FC cutoffs (±1.0, ±0.5, or ±0.25 L_2_FC). These data suggest high ΔΔG values associated with a compound’s ability to regulate full-length LRH-1 in cells, also supported by further analyses in supplemental data. **B**. The same contingency analysis as in A, but replacing ΔΔG with the simple binding energy to full-length LRH-1, comparing compounds in the top 10^th^ percentile of lowest docked binding energies (top 10% or the best 10% binding energies) to full-length LRH-1 *vs*. all other compounds (Remainder), suggesting compounds with lower binding energies did *not* more frequently regulate full-length LRH-1 in cells, at all L_2_FC cutoffs tested, also supported by further analyses in supplemental data. *These data suggest compounds with higher ΔΔG values more frequently regulated full-length LRH-1 in cells*.

### Applying continuous data suggests ΔΔG associates with the ability of compounds to regulate full-length LRH-1 in cells

Our analyses up to this point converted the positive and negative log_2_ fold change continuous data to discrete variables that could be counted, for simplicity. We now plotted the continuous values of *ΔΔ*G for each compound as a function of the absolute value of the log_2_ fold changes *vs*. DMSO control from each of the three wet-lab assays, absolute values were used to account for the equally scaled activation (positive L_2_FC) and repression (negative L_2_FC) by various compounds in the wet lab assays. Spearman rank correlation was significant for the isolated LBD coregulator peptide anisotropy assay (r=0.373, \*\**p*=0.0042) while the slope of linear regression failed to be non-zero by F-test (*p*=0.073, **Fig 7A**). All assays on full-length LRH-1 suggested a significant relationship in both the *CYP17A1*-luciferase data (F-test, \*\**p*=0.0072, Spearman rank r=0.252, \**p*=0.291, **Fig 7B**) and the *CYP8B1-*luciferase data (F-test \**p*=0.018, Spearman r=0.224, \**p*=0.043, **Fig 7C**). Analyzing all full-length LRH-1 assays in cells showed a positive, non-zero slope to the linear regression (F-test \*\*\**p*=0.0005) and a significant Spearman correlation (r=0.399, \*\**p*=0.0010, **Fig 7D**). The positive correlation is being driven by the top 10 of 58 compounds with the highest *ΔΔ*G values, of course during a compound screen these “outlier” compounds with highest activity that are of most interest. To determine if PyRx docking would generate reproducible docking scores (ΔG), as well as ΔΔG values that reproducibly correlate with the cell-based assays, we executed duplicate PyRx docking runs to all 19 structures (Run 2) and directly compared these two independent runs (**Fig 8**). The duplicate runs produced indistinguishable binding scores (**Fig 8A**), ΔΔG values (**Fig 8B**) and correlations with the continuous wet lab data for all cell-based assays (**Fig 8C-E**). These data suggest the PyRx docking scores and ΔΔG reproducibly correlated with regulation of full-length LRH-1 in cell-based assays, between two independent docking runs. We next began to explore these crystal structures more deeply to identify the structural feature being sensed by rigid body docking.

**Figure 7.**
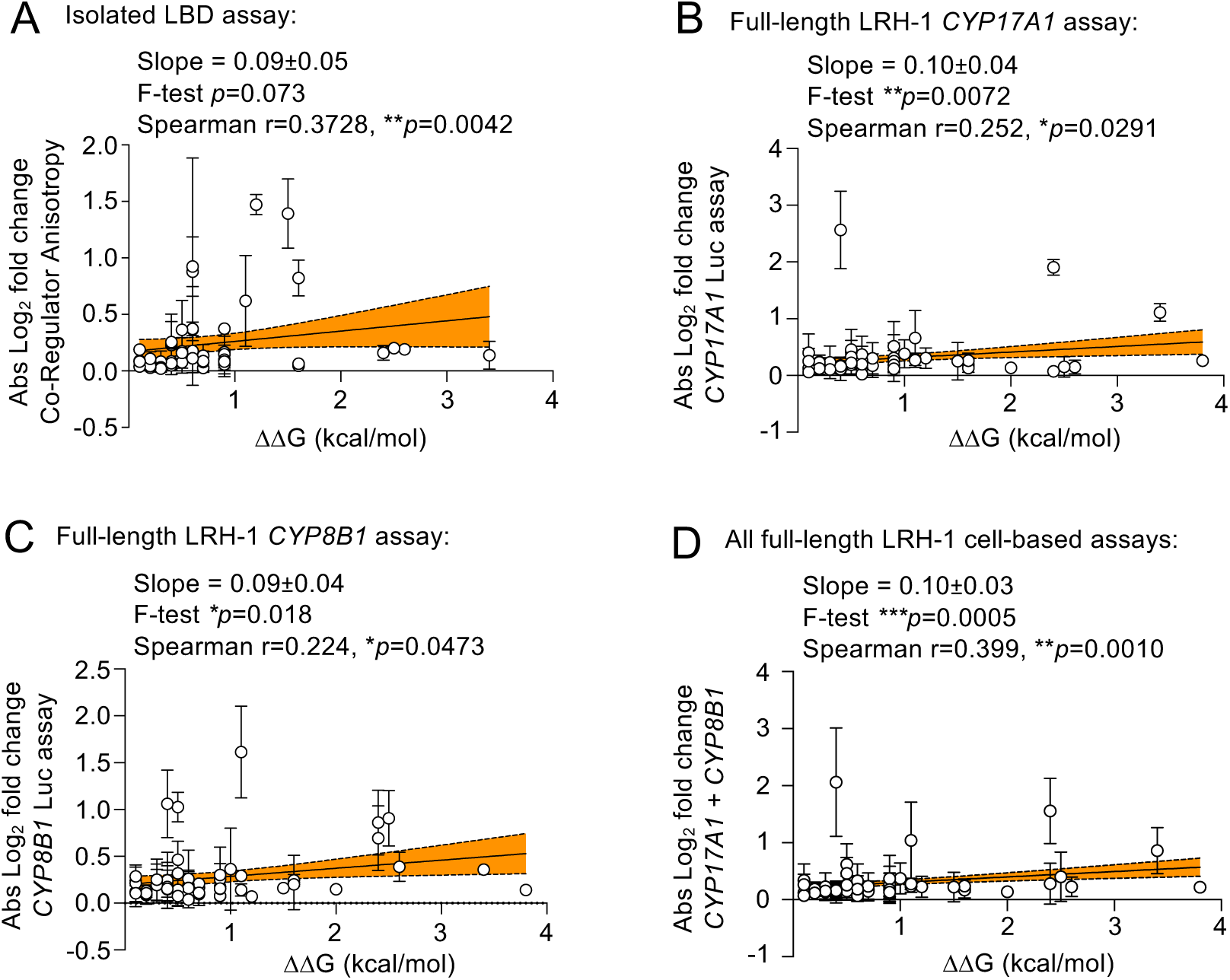
Continuous values of full-length LRH-1 assays positively associate with ΔΔG. Previous analyses converted assay data from the 57 compounds to discrete values since the compounds induced both positive (activation) and negative (repression) log2 fold changes to LRH-1 regulation. Here we plotted ΔΔG values as a function of thecontinous absolute values of the log2 fold change (compared to DMSO control) induced by each compound in each indicated assay: **A.** the co-regulator peptide binding assay using the isolated LBD, **B.** the full-length LRH-1 assay in HEK cells using CYP17A1-luciferase reporter or **C.** the full-length LRH-1 assay in HEK cells using the CYP8B1-luciferase reporter or **D.** all full-length LRH-1 luciferase assays in HEK cells combined. For all plots, solid line indicates linear regression, shaded area is 95% confidence interval, slope and *p* value indicated from F-test for non-zero value of the slope, and the one-tailed Spearman r and *p* values also indicated, calculated in Prism. *These data suggest a positive association exists between ΔΔG of a compound and the ability of that compound to regulate LRH-1*.

**Figure 8.**
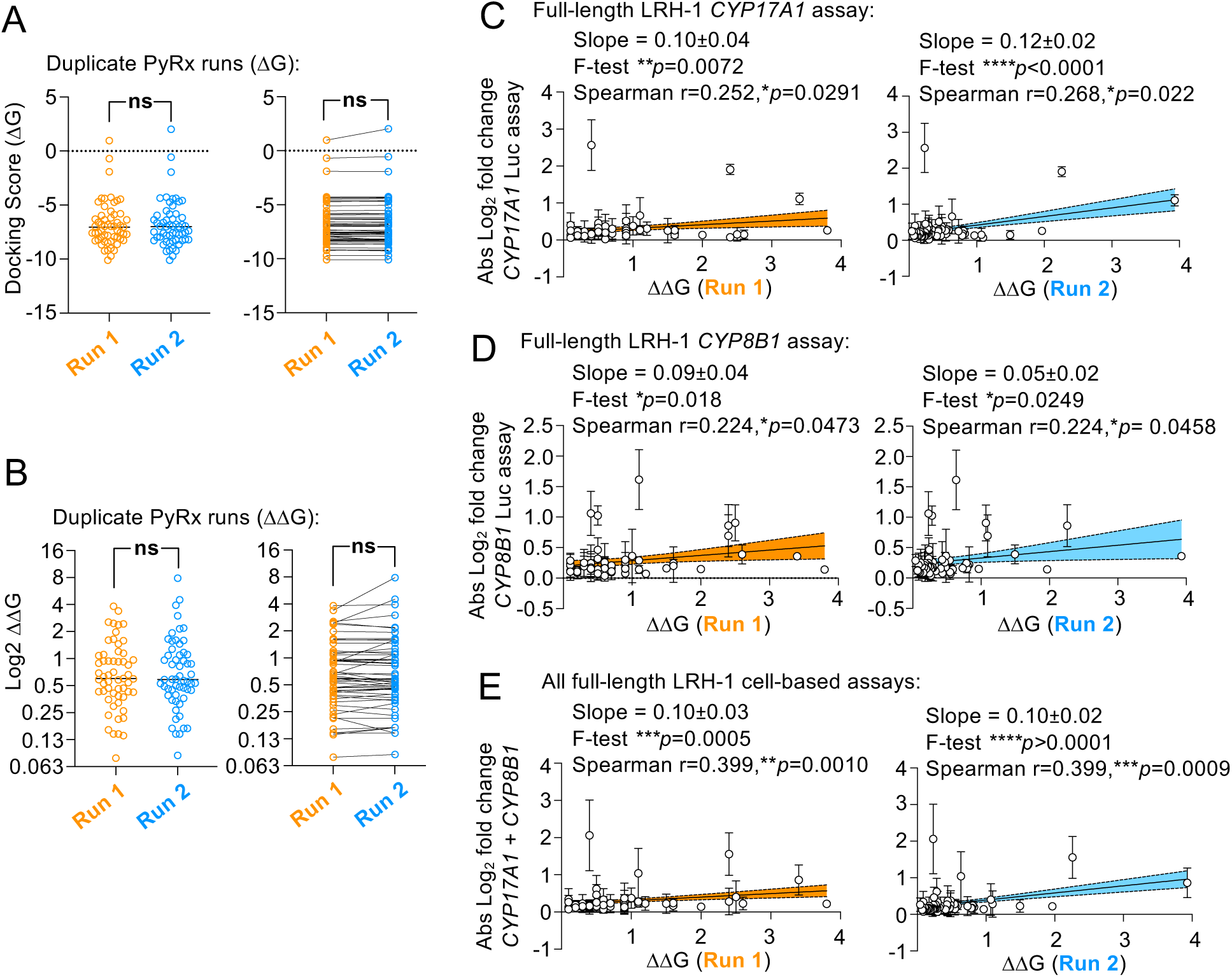
Duplicate PyRx docking runs produced reproducible docking scores (ΔG), ΔΔG values and correlations with LRH-1 regulation. **A**. Comparison of PyRx docking scores (ΔG values) from duplicate runs (Run 1 vs. Run 2) were compared, showing no significant difference by paired t-tests of each of the 57 compounds, each point represents the averaged ΔG from 18 crystal structures of LRH-1. **B**. Comparison of ΔΔG values from duplicate PyRx runs (Run 1 vs. Run 2), again showing no difference in ΔΔG by paired t-tests, suggesting duplicate PyRx runs produced reproducible ΔΔG values, each point represents averaged ΔΔG across 18 crystal structures of LRH-1 for each compound. **C**. ΔΔG values from Run 1 and Run 2 plotted as a function of the absolute value of the log2 fold change (compared to DMSO control) of LRH-1 regulation induced by each compound in the *CYP17A1* cell-based activity assay, showing significant correlation by non-zero linear regression slope and by one-tailed Spearman correlation. **D**. Same as in C. but ΔΔG plotted as a function of CYP8B1 cell-based assays, again showing a significant correlation. **E**. Same as in C, but ΔΔG plotted as a function of both cell-based assays combined. Spearman was one-tailed, all statistics were calculated in Prism. *These data suggest duplicate PyRx docking runs of the 57 compounds to 18 crystal structures of LRH-1 produced reproducible ΔG and ΔΔG values, which correlated with LRH-1 activity in cell-based assays*.

### High ΔΔG values were generated from small-molecule bound LRH-1 LBD crystal structures

Previous work form other groups has established that ligands can induce changes in LRH-1 crystal structures. These ligand-induced differences in co-crystal structures might drive the association of *ΔΔ*G with full-length LRH-1 regulation, therefore the nature of the co-crystalized ligands may indicate how *ΔΔ*G correlates with full-length LRH-1 activity. LRH-1 binds phospholipids, we compared 9 structures in the protein data bank (PDB) of LRH-1 co-crystalized with phospholipids (1YOK, 1YUC, 1ZDU, 3TX7, 4DOR, 4DOS, 4ONI, 4PLE and 4RWV) to another 9 LRH-1 structures co-crystalized with synthetic small molecules (3PLZ, 4PLD, 5UNJ, 5L11, 5SYZ, 6OR1, 6VC2, 6OQX and 6OQY). The *ΔΔ*G values produced by these 18 crystal structures varied for each compound, so we averaged the *ΔΔ*G values for each crystal structure across all 57 compounds, to produce an average *ΔΔ*G value for each crystal structure (*ΔΔ*G_XSTAL_), permitting comparison of crystal structures, rather than comparing compounds. We immediately noted the 9 phospholipid-bound crystal structures all had lower *ΔΔ*G_XSTAL_ values than the 9 small-molecule bound crystal structures (**Fig 10A**), also confirmed in pairwise comparison of the *ΔΔ*G_XSTAL_ values across the 18 structures (**Fig 10B**), consistent with the small-molecule co-crystalized structures producing smaller ligand binding pockets[64]. We next tested if these differences were the source of the association between compound *ΔΔ*G values and full-length LRH-1 regulation by generating a new term *ΔΔ*G* (**Fig 10C**), which is the averaged *Δ*G from the 9 small molecule-bound structures, less the averaged *Δ*G from the 9 phospholipid-bound structures. Although the magnitude of *ΔΔ*G* values were higher than *ΔΔ*G values (**Fig S6A-B**, *p*<0.0001), the value of *ΔΔ*G* did not associate with regulation of full-length LRH-1 in cells, regardless of how the data were analyzed (**Fig 10D**, **Fig S6C**). These data suggest crystal structures that produced the highest *ΔΔ*G values were co-crystalized with synthetic small molecules, while phospholipid-bound structures produced lower *ΔΔ*G values. To identify differences in these two classes of crystal structures, we next applied network analyses.

**Figure 10.**
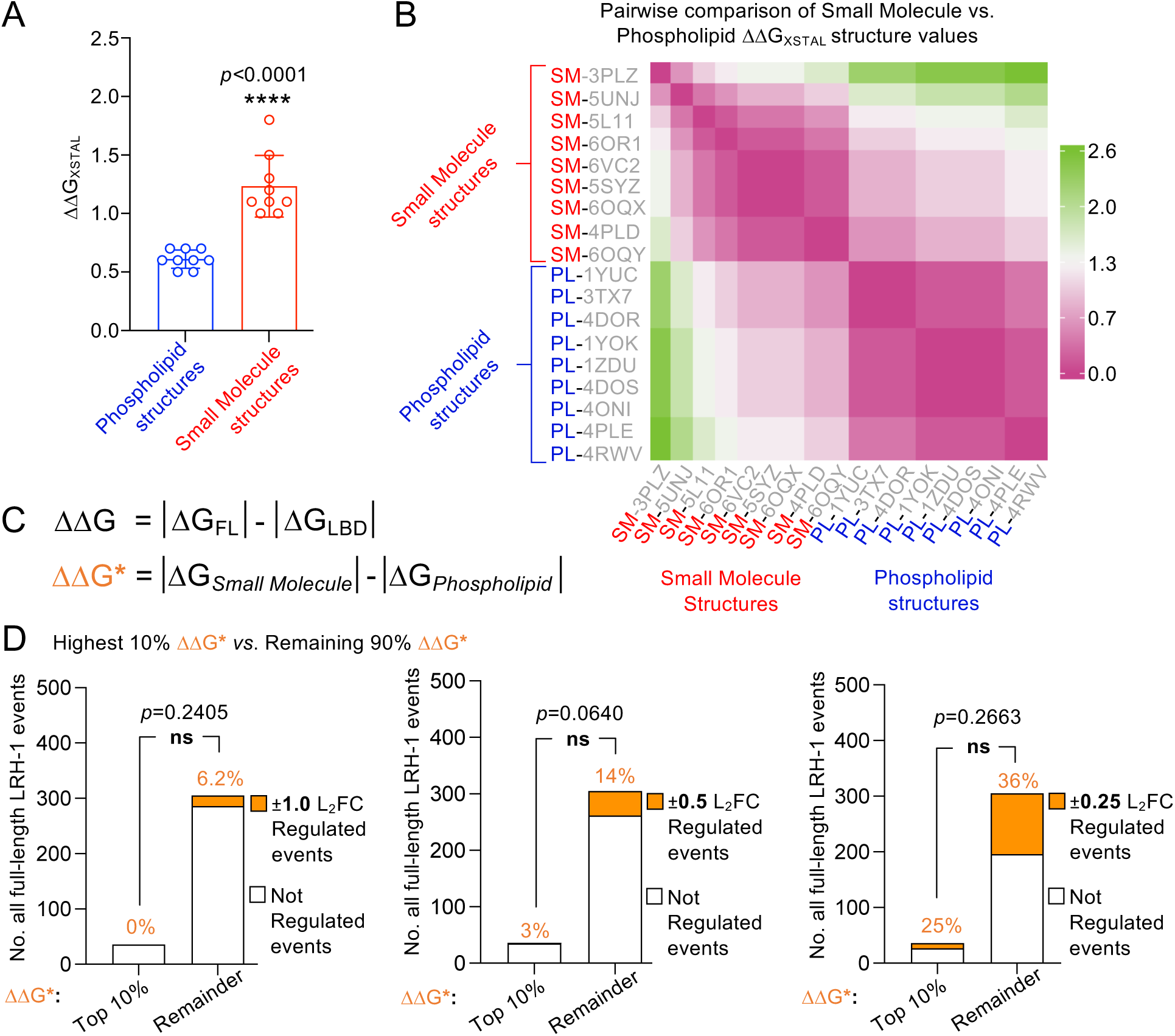
Crystal structures that produced the highest ΔΔG values (ΔΔG_XSTAL_) were bound to small molecules. Since the value of ΔΔG generated from small molecule-bound structures appeared higher than ΔΔG generated from phospholipid-bound structures, we generated a new value called ΔΔG_XSTAL_ which is the average of all 57 ΔΔG values (one ΔΔG value for each of 57 compounds), for each of the 18 crystal structures of LRH-1. **A**. ΔΔG_XSTAL_ for small molecule-bound structures was significantly higher than ΔΔG_XSTAL_ for phospholipid-bound structures (t-test, *p*<0.0001, each point represents ΔΔG_XSTAL_ for one of 18 crystal structures). **B**. Pairwise comparison of all 18 ΔΔG_XSTAL_ values listed by PDB code, suggesting small molecule *vs*. phospholipid-bound structures generated different ΔΔG_XSTAL_ values. **C**. To test if higher ΔΔG values from small molecule-bound structures is the source of the association between *ΔΔ*G and full-length LRH-1 regulation in cells, we determined *ΔΔ*G*, defined as the difference between the average ΔG for each compound bound to the 9 small molecule-bound structures (*Δ*G*_Small Molecule_*) less the average ΔG bound to the 9 phospholipid-bound structures (*Δ*G*_Phospholipid_*). **D.** Contingency showing compounds in the highest 10^th^ percentile of ΔΔG* values did not more frequently regulate LRH-1, regardless of L_2_FC cutoff by Fisher’s exact test, also supported by further analyses in supplemental data, p values calculated in Prism. *These data suggest ΔΔG might reflect a structural aspect of small-molecule bound crystal structures of LRH-1*.

### High ΔΔG_XSTAL_ structures have unique network properties

Several previous studies have used networking approaches to better understand how activating LRH-1 ligands are allosterically paired with transcriptional coregulators[70,71]. To identify any properties of the communication networks within the crystal structures that produced the highest *ΔΔ*G_XSTAL_ values, we performed network analyses on the same 18 structures of LRH-1. We assigned an eigenvector centrality value to each amino acid in each of the 18 structures using RING (3.0)[77] (**Supplemental Spreadsheet 6**). Eigenvector centrality reflects the connectedness of an amino acid with other amino acids in the 3D structure[78]. We then assigned averaged eigenvector centrality across all amino acids in each secondary structural element of the LRH-1 LBD (12 alpha helices, 6 connecting loops and 2 beta sheets, see methods for amino acid numbering, **Supplemental Spreadsheet 6**), and applied principal component analysis to analyze this multi-dimensional data. Variance in the top principal component (PC1) formed two clear clusters: 1) low *ΔΔ*G_XSTAL_ structures bound by phospholipids and 2) high *ΔΔ*G_XSTAL_ structures bound by synthetic small molecules (**Fig 11A**). We then examined edges between nodes in the network (**Supplemental Spreadsheet 7**) that were unique to each of these two groups of crystal structures (**Supplemental Spreadsheet 8**), which showed clear differences between the low *vs.* high *ΔΔ*G_XSTAL_ structures (**Fig 11B**). Superposition of the 18 crystal structures shows that in the nine highest *ΔΔ*G_XSTAL_ structures (**Fig 11C**, **red structures**), Helix 6 is in a position that constricts the entrance to the ligand binding pocket (**Fig 11D**), also suggested by the position of Helix 3 closer to Helix 6 (**Fig 11E**). In the nine lowest *ΔΔ*G_XSTAL_ structures Helix 6 is in the opposite position, opened at the entrance to the ligand binding pocket (**Fig 11C-E**). Previous network studies based on molecular dynamics simulations of the ligand binding domain have also suggested the position of Helix 6 as an important determinant of LRH-1 conformations that associate with ligand regulation[71]. The data presented here now suggest the position of Helix 6 correlates with *ΔΔ*G values, and *ΔΔ*G correlates with full-length LRH-1 activity in cell-based assays. The data here suggest *ΔΔ*G has utility in prioritizing hit compounds from screening campaigns for follow-up screening in full-length LRH-1 cell-based assays.

**Figure 11.**
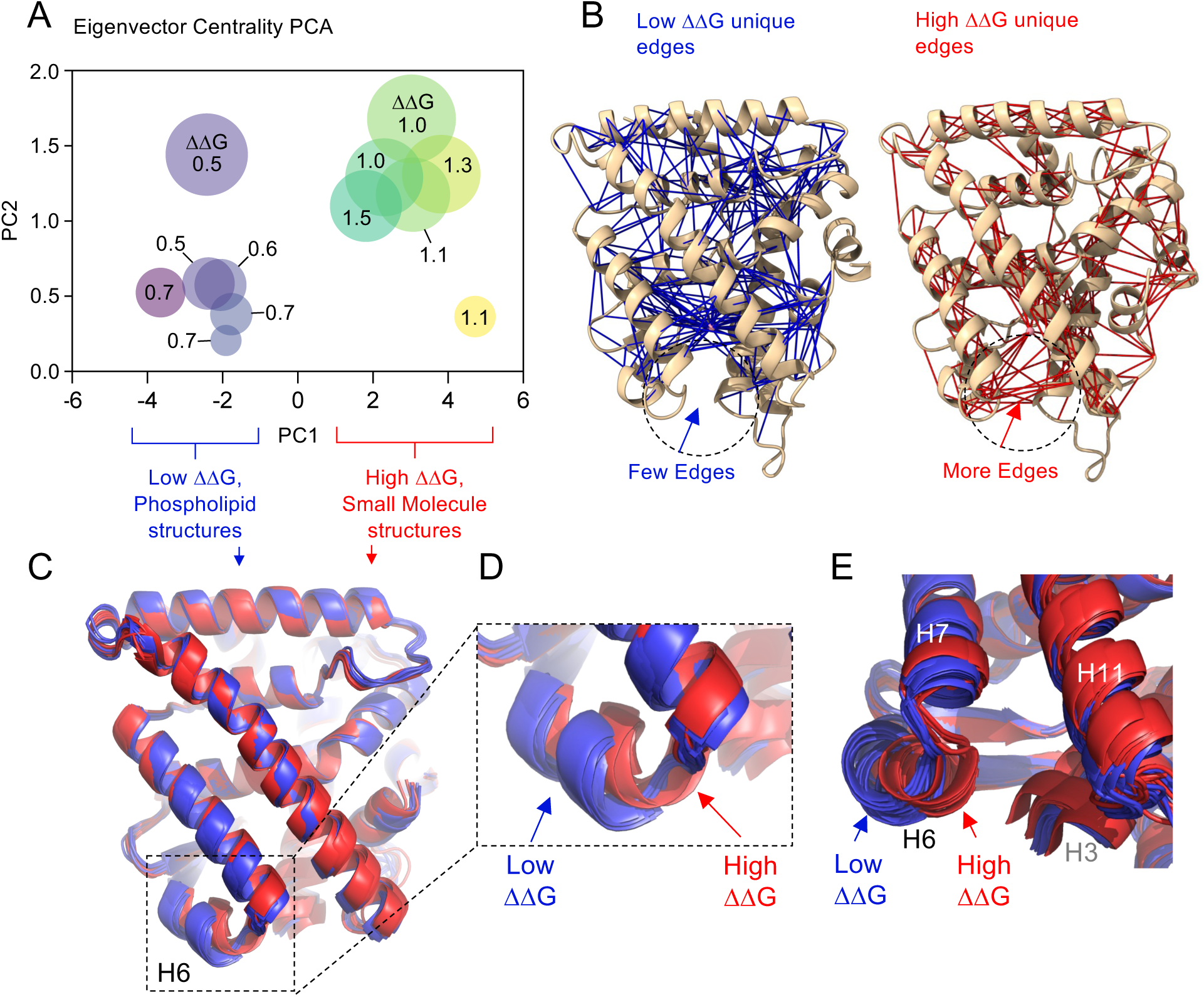
The position of Helix 6 associates with higher ΔΔG_XSTAL_ values. The ΔΔG_XSTAL_ is the averaged ΔΔG value of 57 compounds for each crystal structure (indicated in this figure simply as ΔΔG). **A.** Principal Component Analysis of eigenvector centrality values assigned to each secondary structural element (12 Helices, 3 loops and 1 beta strand) in all 18 crystal structures of LRH-1. PCA shows clustering of structures with low ΔΔG_XSTAL_ values co-crystalized with phospholipids *vs.* structures with high ΔΔG_XSTAL_ values co-crystalized with small molecules, see methods for network analysis details, ΔΔG_XSTAL_ values indicated. **B.** Edges between network nodes that were unique to the low ΔΔG_XSTAL_ structures (blue, left) or the high ΔΔG_XSTAL_ structures (red, right) were mapped onto PDB:6OQX. Note that among many differences, one difference in the high ΔΔG_XSTAL_ structures is the presence of more edges between Helix 6 and Helix 3 at the entrance to the ligand binding pocket, indicated with dashed circle and arrows. **C.** Superposition of all 9 low ΔΔG_XSTAL_ structures bound by phospholipids (blue) and all 9 high ΔΔG_XSTAL_ structures bound by small molecules (red), less ordered loop regions were removed for clarity. Examination of the superposition showed the largest change in position was associated with Helix 6. **D**. Closeup of boxed region in panel C., showing change in Helix 6 position in the low ΔΔG_XSTAL_ vs. high ΔΔG_XSTAL_ structures. **E**. End-on alternative view of Helix 6, showing the position of Helix 6 favoring closer proximity to Helix 3 in the high ΔΔG_XSTAL_ structures. *These data suggest that positions of Helix 6 which close the entrance to the ligand binding pocket associate with higher values of ΔΔG*.

## Discussion

Prioritizing hit compounds from a large screen is often a necessary experimental constraint[1–6] as secondary assays in living cells can be very resource-intensive[7,8,13]. Therefore in terms of drug screening, the few “outlier” compounds with the most biological activity are of the highest biomedical interest. Identifying these high activity “outlier” compounds is the goal of drug screening. Here, we present a computational heuristic (*ΔΔ*G) with utility prioritizing LRH-1 compounds for secondary assays in cells. We validated the *ΔΔ*G metric retrospectively by using a total of 439 orthogonal functional assays. We also present data suggesting the more direct metric of docked binding energy to LRH-1 did *not* correlate with the ability of compounds to regulate full length LRH-1 in cells. This study identifies and validates a link between *in silico* docking score and LRH-1 function in cell-based high throughput functional assays. Binding constants determined in the wet lab have been linked to LRH-1 regulatory activity for individual compounds[42,49], however these approaches are challenging and expensive to scale up. We propose *ΔΔ*G could be applied to prioritize hits from large primary compound screens targeting LRH-1.

Although it is difficult for the computational data presented here to definitively establish how *ΔΔ*G associates with LRH-1 regulation, we found three lines of evidence suggesting a connection with the position of Helix 6 at the entrance to the LRH-1 ligand binding pocket. First, the docked positions of high *ΔΔ*G compounds clustered around the entrance to the ligand binding pocket at Helix 6 (**Fig 2**), suggesting changes to the position of Helix 6 might be responsible for altered compound docking and higher *ΔΔ*G values. Second, our network analyses suggested the crystal structures that generated the highest *ΔΔ*G_XSTAL_ values had more edges connecting Helix 6 to Helix 3. Third, the entrance to the ligand binding pocket in high *ΔΔ*G_XSTAL_ structures was in a more closed position. Here we must mention that comparing crystal structures from non-identical space groups must be interpreted with caution. Perhaps most importantly, convincing work from the Ortlund lab at Emory and now the Okafor lab at Penn State[74] have used extensive MD simulations, network analyses and direct comparisons of different ligand-bound structures of LRH-1 to independently suggest an important role for Helix 6 in ligand-regulated activation of LRH-1[37,64,70,71,74–76], linking the mobility of residues at the entrance to the ligand binding pocket near Helix 6 to synthetic small molecule-induced LRH-1 activity[76]. The volume of the ligand-binding pocket is smaller in structures bound to synthetic small molecules vs. phospholipids[64], consistent with the docking studies presented here, as well as hydrogen-deuterium exchange data[37,64,71,76], MD simulations[64,70,71,74–76] and other network analyses[70,76]. Together, these studies support a role for Helix 6 is an important element in translating ligand binding events from the binding pocket to the transcriptional coregulator[70,71]. Our analyses of cell-based data are consistent with those findings, however it will be important to apply structural biology to determine how high *vs*. low *ΔΔ*G compounds regulate the dynamics of Helix 6 in future analyses.

The structures generating the highest *ΔΔ*G values were co-crystalized with synthetic small molecules, while the phospholipid-bound structures produced the lowest *ΔΔ*G values. The full-length LRH-1 structure is a computational model based on a phospholipid-bound LBD crystal structure (PDB:1YOK). We therefore hypothesized it might be this phospholipid-bound starting point in the full-length modeling that produced high *ΔΔ*G values. Calculating *ΔΔ*G* using the average binding energies of each compound to the phospholipid-bound *vs.* synthetic small molecule-bound LBD crystal structures produced *ΔΔ*G* values that did not associate with any regulation of full-length LRH-1 in cells. It therefore remains unclear what aspects of rigid docking to the full-length LRH-1 results in the association between *ΔΔ*G and full-length LRH-1 regulation in cells, but we will speculate on two potential hypotheses. The first is rather trivial, simply that the modeled position of Helix 6 in full-length LRH-1 is somewhat “between” the position of Helix 6 in the high *vs*. low *ΔΔ*G_XSTAL_ structures, and that this position imparts the ability of the full-length model to uniquely “sense” or “block” interactions between compounds and LRH-1. The second is highly speculative but worth mentioning. The published model of full-length LRH-1 does not position Helix 6 in the interface between the LBD and the DNA-binding domain (DBD). However, while developing the full-length LRH-1 model[73], a lower-energy model was produced by Rosetta, in which Helix 6 was directly in the LBD-DBD interface, as were residues at the entrance to the ligand binding pocket, which we referred to as “Model 2”. Although Model 2 was supported by certain biophysical experiments (specifically chemical crosslinking), Model 2 was genetically tested using mutants of LRH-1 (I415Q and S418D) which did not affect full-length LRH-1 activity[73], eliminating Model 2 from further consideration. Still, it remains formally possible that in a cellular context that remains untested, Helix 6 could exist in the interface between the LBD and DBD in particular conformational states of full-length LRH-1. Accordingly, the high *ΔΔ*G compounds might simply bind LRH-1 at the entrance to the ligand binding pocket near Helix 6 (**Fig 2E**), directly in the LBD-DBD interface predicted by Model 2. Supporting this hypothesis, compounds that operate in this way would be expected to bind the isolated LBD (**Fig 1B**)[1] and regulate full-length LRH-1 (**Fig 1G**) but would not be expected to regulate the isolated LBD (**Fig 1F**). A total of 9 compounds of the 57 examined here meet those criteria (**Supplemental Spreadsheet 2**) and will be of great interest to determine co-crystal structures. More wet-lab study will be needed to establish the role of Helix 6 in full-length LRH-1 structural regulation.

One limitation of this study is the small size of the compound library (only 57 compounds from a 2322 compound library), as it remains possible that in larger (or smaller) screens the *ΔΔ*G metric would not associate with compound activity on full-length LRH-1 in cells. However, the small size of the primary screen permitted application of several independent, orthogonal secondary assays, a clear strength of this study. Executing more compound screens using different library sizes with follow-up secondary assays in cells would be a very resource-intensive task and outside the scope here. The secondary screens in our previous paper were limited to only the hit compounds identified, we did not perform secondary screening on all 2322 compounds, as resources are not unlimited. That our analyses found correlations in the data suggests similar associations might hold in larger screens, which can improve ongoing LRH-1 compound screening and development efforts. Thus, the data here can have practical impact on LRH-1 drug screening. Regardless of how ΔΔG associates with cell-based LRH-1 activity. the utility of ΔΔG in prioritizing hits from wet lab drug screens remains.

This study introduces a new metric called *ΔΔ*G, derived from the rigid docking scores of compounds docked to 19 different structural models of LRH-1, which positively correlates with the ability of 57 compounds to regulate full-length LRH-1 in cell-based assays. The docking scores themselves did not have any association with full-length LRH-1 activity in cells. Network analyses suggest a closed position of Helix 6 at the mouth of the ligand binding pocket associates with the highest *ΔΔ*G_XSTAL_ structures, also observed in other computational studies of LRH-1[71]. This observation provides a potential explanation as to how *ΔΔ*G associates with compound activity on LRH-1, which awaits further testing using wet lab structural biology. This study provides a new computational tool that can aid in the prioritization of hit compounds for follow-up secondary screens in LRH-1 drug development efforts, while further supporting an important role for Helix 6 in ligand-regulation of full-length LRH-1.

## Methods

### Materials

All new data in this manuscript is computational, all wet lab assay data were previously published[1], with the most relevant data provided as Supplemental Spreadsheets with this manuscript for convenience. Rigid body docking was executed with PyRx (Version 0.8) using Autodock Vina[79], run on a [79]single Dell Precision 5820 machine running Ubuntu Linux 20.04 on an Intel Xeon 8-core, 3.9GHz processor. Structures were visualized and figures generated using Schrodinger PyMOL (Version 2.4.2) and UCSF ChimeraX (Version 1.6.1), protein structure networks were generated using RING (Version 3.0) mapped onto structures by UCSF ChimeraX, eigenvector centrality was calculated using python library NetworkX (Version 3.1) and data were processed in Microsoft Excel (Version 16.74).

### Statistical analyses

Principal component analyses used equally scaled data for all variables analyzed (log2 fold change compared to DMSO control) and parallel analysis with 1000 simulations and selected the 2 components shown in biplots in Figure 1 and Figure 11 were calculated in GraphPad Prism (Version 10.0.0), the matrices for the PCA in Fig 1E is provided as Supplemental Spreadsheet 1, and for Figure 11A the PCA matrix is provided in Supplemental Spreadsheet 6. Fisher’s exact tests of contingency were two-sided for all analyses also calculated in Prism. Spearman rank correlation r values were approximated, all Spearman *p* values were one-tailed as the slope of initial linear regressions were positive making negative correlations unlikely. Linear regressions, slopes and F-tests for non-zero slopes were also calculated in Prism, regressions were performed without constraint on x-axis intersection. Pairwise comparison of ΔΔG_XSTAL_ values used a Euclidean distance matrix calculated by Heatmapper.ca, paired t-test was two-tailed and calculated in Prism, all analysis files are available upon request. The statistical tests are listed in figure legend, “ns” represents any p value that is not significant (tests null) for indicated test, “ *p” represents any p-value less than 0.05, “ **p” represents any p-value less than 0.01, “ ***p” represents any p-value less than 0.005, “ ****p” represents any p-value less 0.0001. All analysis files are available on request.

### Exclusion of VU0656093

Although 58 compounds were identified in the previously published compound screen, only 57 compounds were analyzed for the majority of this study. We excluded one compound (VU0656093) based on the PyRx docking to human LRH-1 ligand binding domain crystal structure PDB:6OQX from our previously published data[1], as PyRx docking resulted in an extraordinarily high positive binding energy (+44.9 kcal/mol) to 6OQX. In this study, VU0656093 was also the only compound to produce positive values for the docked binding energy to the LBD structure 1YOK (+15.3kcal/mol) and the full-length LRH-1 model (+26.6kcal/mol). PyRx relative docked binding energies for VU0656093 are included in **Supplemental Spreadsheet 3** associated with this manuscript for completeness.

### Overview of published wet lab data

The 58 hit compounds were identified in a previous publication to directly bind the isolated LBD of LRH-1, using a FRET-based high-throughput screen[1]. Briefly, the 2322 compound Discovery Spectrum Collection library was used in that FRET screen, the 58 compounds were identified to decrease FRET between a fluorophore-labeled phospholipid probe installed in LRH-1 at the canonical ligand binding site and the fluorophore-labeled LRH-1 LBD protein, as FRET donor and acceptor respectively. The design of the screen suggests the identified hit compounds compete with the phospholipid probe for binding to the canonical ligand binding site in LRH-1, however no structures of the 58 hit compounds co-crystalized with LRH-1 have demonstrated this unequivocally. Binding constants to LRH-1 were not determined for most hit compounds from the screen, but a handful of binding constants were determined to validate the screen, including two high *ΔΔ*G compounds relevant to the current study, VU0243218 (IC_50_=9.4μM [95%CI 8.1-10.8μM] for binding to the isolated LRH-1 LBD) and VU0656021 (IC_50_=27.0μM [95%CI 16.1-171.5μM] for binding to the isolated LRH-1 LBD). Both these compounds are high *ΔΔ*G compounds that had fully saturable binding curves that could only be fit to a one-site binding model, strongly suggesting a direct, stoichiometric interaction of at these two compounds with the LRH-1 ligand binding domain, despite docking outside the canonical ligand binding pocket of LRH-1 (PDB:6OQX). The 58 hit compounds were subjected to secondary functional screens: 1) 10μM compound-induced interaction between recombinantly expressed and purified LRH-1 LBD and a fluorophore-labeled peptide (representing the transcriptional coactivator PGC1α) by fluorescence anisotropy. 2) 10μM compound-induced luciferase expression of a *CYP8B1*-promoter driven reporter in HEK293T cells expressing full-length human LRH-1 by transient co-transfection. Importantly, control luciferase reactions in the absence of co-transfected LRH-1 were used for normalization, so only the compound-regulated luciferase signal dependent upon LRH-1 co-transfection was analyzed. 3) 10μM compound-induced luciferase expression of a *CYP17A1*-promoter driven reporter in HEK293T cells expressing full-length human LRH-1 by transient co-transfection, using the same control methods as above. Results from the previously published assays are available associated with this manuscript in Supplemental Spreadsheet 1, for complete technical details on all the methods and data see DOI: 10.1021/acschembio.2c00805.

### PyRx computational docking

PyRx[79] rigid body computational docking was used, the 58 hit compounds were first docked to 6OQX PDB structure of human LRH-1 ligand-binding domain (LBD)[1], then docked to the integrated structural model of full-length human LRH-1 (PDB_DEV: 00000035), then PDB 1YOK as another crystal structure of human LRH-1 LBD, followed by the remaining 16 crystal structures analyzed in this study for a total of 18 human LRH-1 LBD crystal structures used for PyRx docking, listed here (PDB: 6OQX, 6VC2, 6OR1, 6OQY, 5UNJ, 5SYZ, 5L11, 4RWV, 4PLE, 4PLD, 4ONI, 4DOR, 4DOS, 3TX7, 3PLZ, 1ZDU, 1YUC, 1YOK), plus the integrated structural model of full-length LRH-1 (PDB_DEV: 00000035). The full-length LRH-1 structure is based on in-solution biophysical restraints applied to Rosetta-based computational docking, in which the ligand binding domain was computationally optimized, which was then validated using genetics, biochemistry and solution structural analyses[73] but is not a crystal structure. For PyRx ligand docking, all proteins were prepared for docking by removing all co-crystallized ligands, ions, and water molecules. The 2D ligand structures in SDF format were converted to 3D using OpenBabel[80] and energy-minimized using a universal force field with 200 steps, saved in PDBQT format, all ligands are available in Supplemental_Zip_File_1.zip available on the publisher’s website or by contact the corresponding author. Size and position of the grid box, search space, and scoring function were set in PyRx[72], the docking grid box size was X:25Å, Y:25.2Å, Z:25Å centered on the canonical ligand binding site in human LRH-1. PyRx generated nine docked poses for each ligand with a corresponding docking score (relative docked binding energy, kcal/mol), pose associated with the lowest energy docking score was used for analyses. Output files were saved in PDBQT format, all docked poses were visualized using molecular graphics software PyMOL[81] or UCSF ChimeraX[82]. Docking scores are provided in Supplemental Spreadsheet 3, PDB files for all docked poses are available upon request and are included in the Supplemental_Zip_File_1.zip.

### Protein structure networks

Connectivity within sets of LRH-1 crystal structures were evaluated using protein structure networks (PSN) generated using RING 3.0[77]. Identical parameters were selected for all 18 structures (PDB: 6OQX, 6VC2, 6OR1, 6OQY, 5UNJ, 5SYZ, 5L11, 4RWV, 4PLE, 4PLD, 4ONI, 4DOR, 4DOS, 3TX7, 3PLZ, 1ZDU, 1YUC, 1YOK), using a relaxed model that returns one edge between two amino acid nodes. Protein structure networks in RING were set to include the closest nodes using relaxed distance thresholds, a single edge, with water molecules excluded and distance parameters set to 5.5Å for hydrogen bonds, 5Å for ionic interactions, 0.8Å for Van der Waals, 7Å for ρε-ρε stacking, 7Å for ρε-cation interactions and 3Å for disulfide bonds. Protein structure networks were generated using edge files by creating an adjacency list from Nodes *a* and *b*, each representing the two structures being pairwise compared (*a*,*b*), and the edges mapped between the C-alpha atom coordinates from each PDB file. The network edges were imposed on protein structures for visualization by mapping pseudo bond structures in UCSF ChimeraX[82], the NetworkX Python package was used to generate the graph objects [77][82].

### Eigenvector centrality and principal components analysis

Protein structure networks generated by RING 3.0[77] were used in eigenvector centrality (EC) analysis using python library NetworkX to yield an EC value for each amino acid node in the network for 18 LRH-1 crystal structures from the PDB (PDB: 6OQX, 6VC2, 6OR1, 6OQY, 5UNJ, 5SYZ, 5L11, 4RWV, 4PLE, 4PLD, 4ONI, 4DOR, 4DOS, 3TX7, 3PLZ, 1ZDU, 1YUC, 1YOK), the same 18 crystal structures used throughout this study. EC values are reported in **Supplemental Spreadsheet 5**. [77]EC values at amino acid nodes were averaged within the standard secondary structural elements of the LRH-1 LBD, each element assigned a label according to the standard 12-helix ligand binding domain as Helix 1(300-310), Helix 2(314-330), Loop 2(331-339), Helix 3(340-362), Helix 4(365-369), Helix 5(370-397), Loop 4(398-401), Helix 6(413-418), Helix 7(421-442), Helix 8(444-457), Loop 8(458-465), Helix 9(466-489), Loop 9(490-493), Helix 10(494-501), Helix 11(502-523), Loop 10(524-529), and Helix 12(530-538). For each of these 17 secondary structural elements, the EC values of all amino acid nodes in the element were averaged, averaged EC values for each secondary structural element in LRH-1 were used as 17 features for principal component analyses. PCA used 1000 simulations, executed in GraphPad Prism (Version 10.0.0). Eigenvector centrality (EC)is a metric of how strongly the centrality score of a node in a network is influenced by its connections to other central nodes. The largest eigenvalue of the adjacency matrix scales the magnitude of the associated eigenvector and satisfies the equation Av=λv, where A is the network adjacency matrix and λ is the largest eigenvalue of the adjacency matrix. This implies that for a network of size j, the eigenvector value for a given vertex v_i_ is such that v_i_ = λ^-1^ ∑_j_A_ij_ v_j_. In other words, the eigenvector value v_i_ is the sum of the eigenvector values of its first-degree neighbors scaled by the largest eigenvalue.

### Identifying unique network edges

To identify edges unique to each cluster in the principal component analysis, all unique edges for structures with high *ΔΔ*G values were grouped together in Excel (PDB: 3PLX, 4IS8, 4PLD, 5I11, 5SYZ, 5UNJ, 6OQX, 6OQY, 6OR1, 6VC2) and all unique edges for structures with low *ΔΔ*G values grouped together (PDB: 1YOK, 1YUC, 1ZDU, 3TX7, 4DOR, 4DOS, 4ONI, 4PLE, 4RWV). Common edges between the two groups were identified and removed in Excel, edges outside the intersection for each cluster were mapped onto PDB:6OQX by mapping pseudo bond structures using UCSF ChimeraX[82].

## List of supplemental spreadsheets

**Supplemental Spreadsheet 1**: Published continuous wet lab assay data[1].

**Supplemental Spreadsheet 2**: Published functional assays from 14 hit compounds[1].

**Supplemental Spreadsheet 3**: PyRx relative docked binding energies to LRH-1.

**Supplemental Spreadsheet 4**: Matrix of all 1026 *ΔΔ*G values.

**Supplemental Spreadsheet 5**: Eigenvector centrality values for 18 LRH-1 structures.

**Supplemental Spreadsheet 6**: Average Eigenvector centrality across LRH-1 secondary structures.

**Supplemental Spreadsheet 7**: All network edges.

**Supplemental Spreadsheet 8**: Unique edges to high vs. low *ΔΔ*G_XSTAL_ structures.

**Supplemental Spreadsheet 9**: PyRx Run 2 relative docked binding energies to LRH-1.

## Data availability

All results associated with this manuscript are provided as supplemental spreadsheets, all other files are available upon request.

---

**Supplemental Fig. S1.**
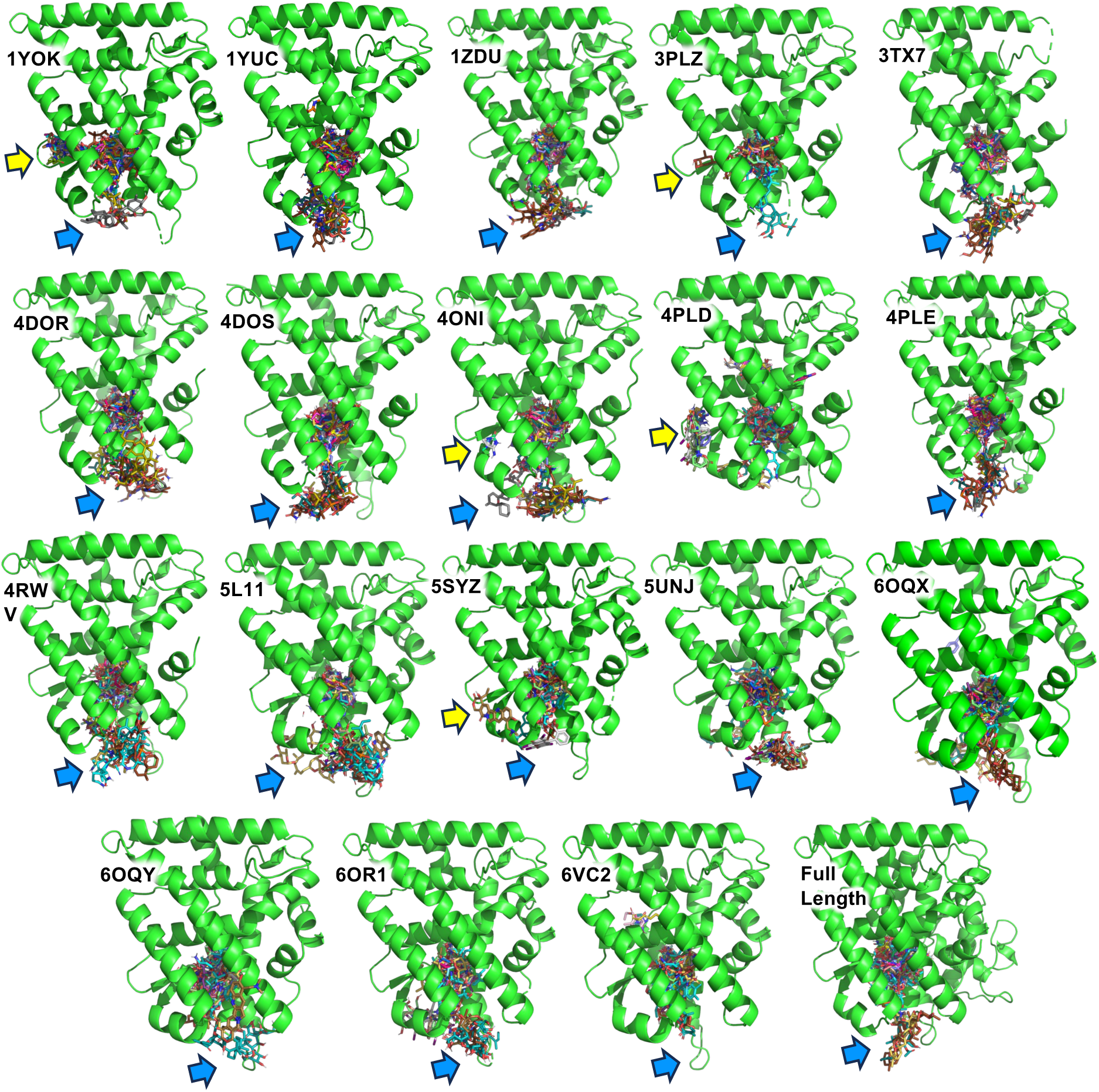
Most of the 57 compounds docked to either the canonical ligand binding site or the mouth of the ligand binding pocket in LRH-1. Indicated PDB crystal structures (green ribbon) used for rigid PyRx docking to the 57 hit compounds (colored sticks) to illustrate the sites where compound docking occurred, 3 sites were identified. Site 1 is not labeled but is the canonical ligand binding site, where the majority of compounds docked, in the central core of the LRH-1 LBD. A second non-canonical site (**blue arrows**) was near Helix 6 at the mouth of the ligand binding pocket. Docking at this site is consistent with published studies suggesting ligand binding to this region is important for ligand regulation of LRH-1. A third non-canonical site (**yellow arrows**) was observed in a few structures, particularly 4PLD which represents the apo-state of LRH-1. *These data suggest the majority of compounds docked to the canonical ligand biding site and to a non-canonical site at the mouth of the ligand binding pocket in human LRH-1*.

**Figure S2.**
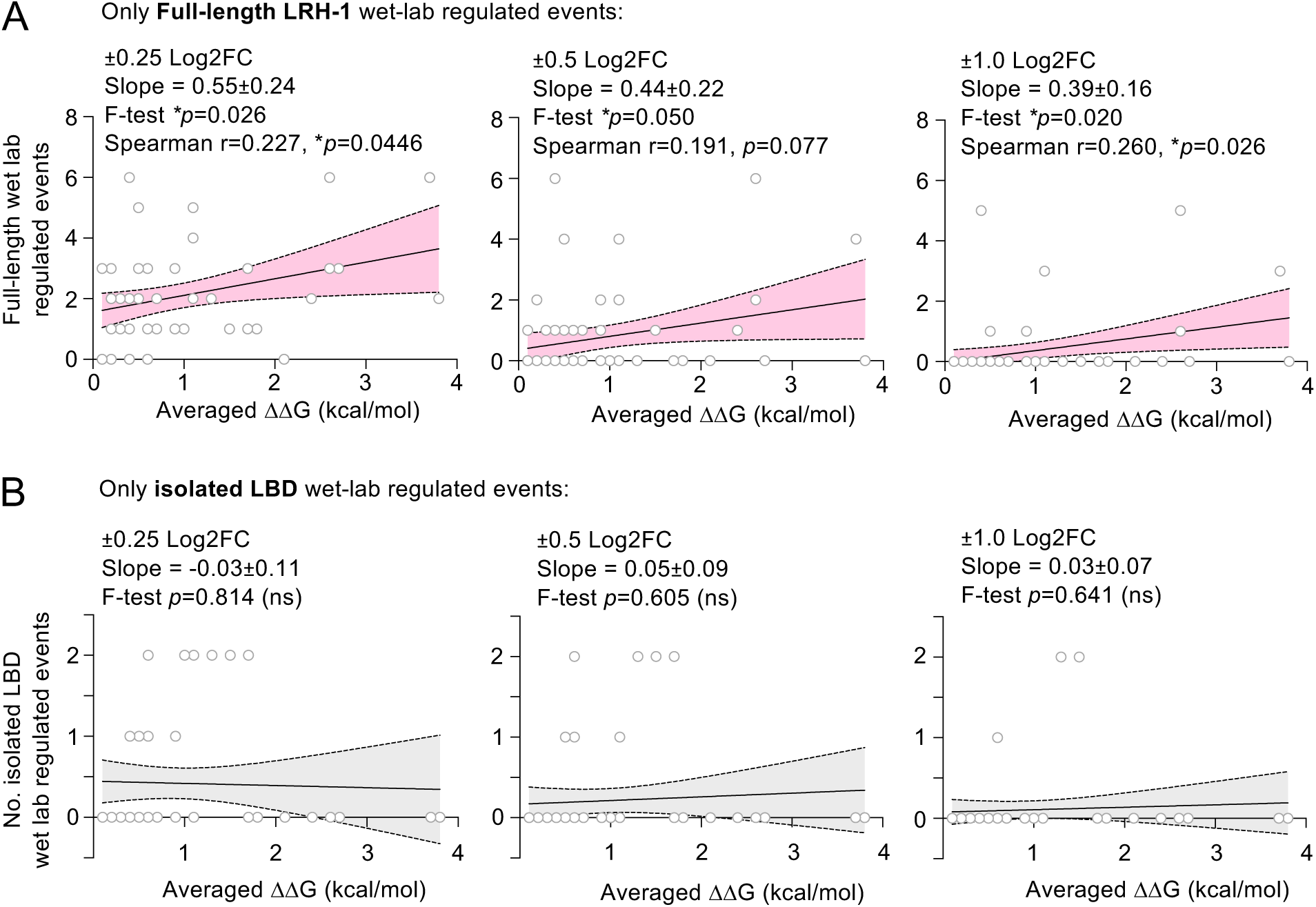
A positive correlation exists between ΔΔG averaged across 18 crystal structures of LRH-1 and the number of compound-regulated full-length LRH-1 events in cells. **A**. 57 of 58 hit compounds (one compound was excluded, VU0656093, due to high positive energies docked to several LRH-1 structures) were docked to 18 crystal structures of the human LRH-1 ligand binding domain (LBD) and ΔΔG values calculated, producing a matrix of 1026 ΔΔG values, provided in supplemental data. The average of the 18 ΔΔG values across the 18 crystal structures examined for each of the 57 compounds was plotted as a function of the number of all LRH-1 wet-lab regulated events that were induced by each compound, with regulation defined as at least ±0.25 Log2 fold change (L_2_FC) from DMSO control (left plot), ±0.50 L_2_FC (middle plot) or ±1.0 L_2_FC (right lot). **B**. Identical as in A., except that only isolated LBD wet lab events were plotted against the same ΔΔG values, failing to show a significantly non-zero slope. For all plots, solid line indicates linear regression for all points, shaded area is the 95% confidence interval for the regression, slope of the regression and *p* value from F-test for a non-zero slope are indicated. *These data suggest a relationship between ΔΔG and a compound’s ability to regulate full-length LRH-1 in cells, warranting further investigation*.

**Figure S3.**
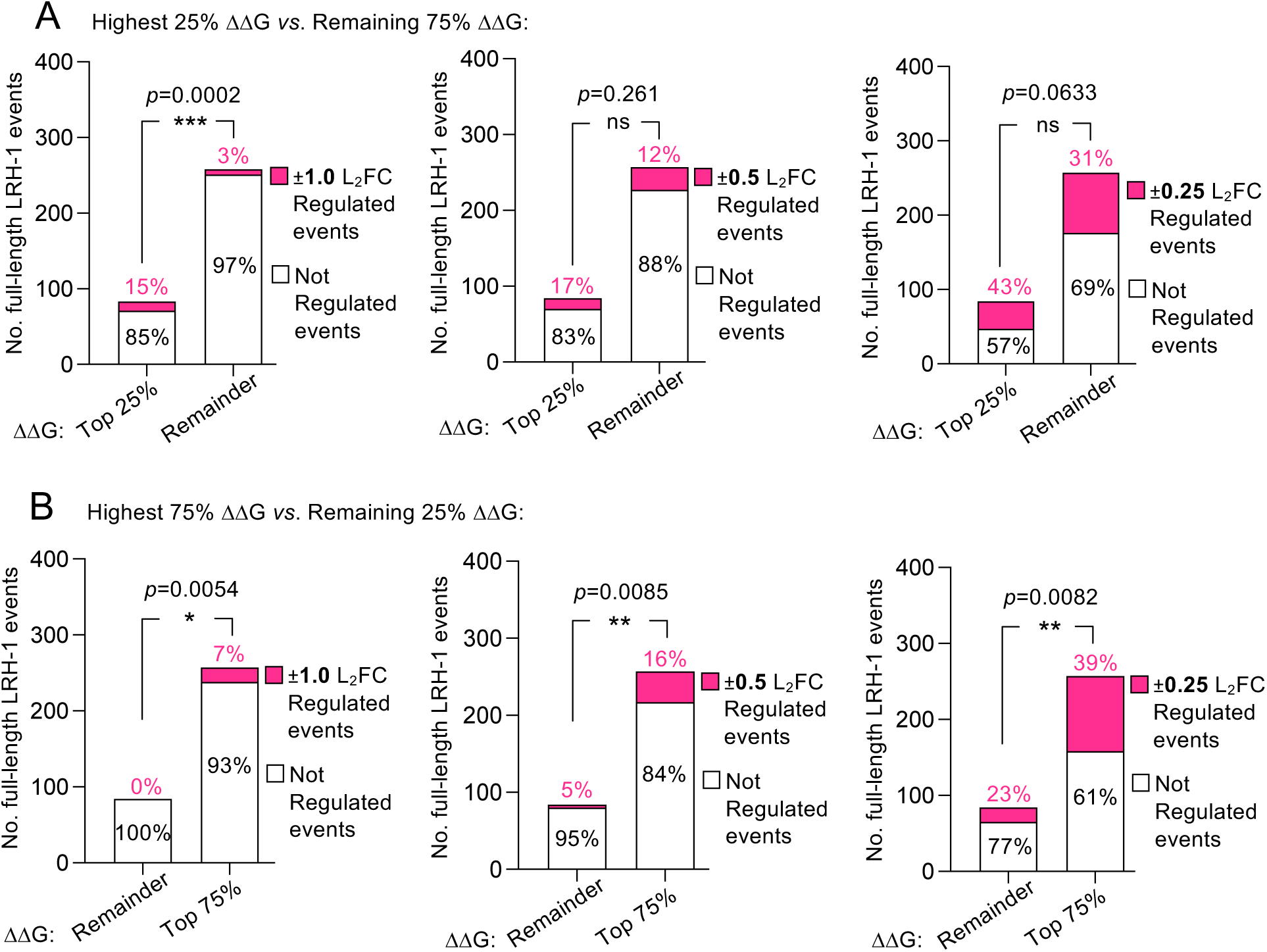
Compounds with high values of ΔΔG associate with regulation of full-length LRH-1 in cells. Contingency analyses of the frequency of LRH-1 regulation induced by all 57 compounds in all 341 full-length LRH-1 assayed events in cells (y-axis, each of the 341 wells in the high-throughput assay treated as an individual event, i.e. replicates not averaged) and regulation defined as meeting the indicated minimum log_2_ fold change cutoff (L_2_FC) compared to DMSO control. **Pink section** of bars is the number of regulated events (percentages indicate percentage of events that were regulated), **white section** of bars is the number of unregulated wet lab assayed events, e.g. that did not meet the minimum L_2_FC cutoff. **A**. Contingency analysis of the frequency of LRH-1 regulation in all 341 assays, comparing compounds in the top quartile of ΔΔG values *vs*. all other compounds, showing compounds with higher ΔΔG values more frequently regulated LRH-1 at ±1.0 L_2_FC (15% vs. 3%, *p*=0.0002), while the less stringent ±0.5 and ±0.25 L_2_FC did not reach statistical significance. **B.** Contingency analysis of the frequency of LRH-1 regulation in all 341 assays, comparing compounds in the top 3 quartiles of ΔΔG values (Top75%) vs. all other compounds (Remainder), showing compounds with higher ΔΔG values more frequently regulated LRH-1 at ±1.0 L_2_FC (7% vs. 0%, *p*=0.0054), at ±0.5 L_2_FC (16% vs. 5%, *p*=0.0085) and the most inclusive ±0.25 L_2_FC (39% vs. 23%, *p*=0.0082). *These data suggest the value of ΔΔG positively associates with the ability of a compound to regulate full length LRH-1 in cells*.

**Figure S4.**
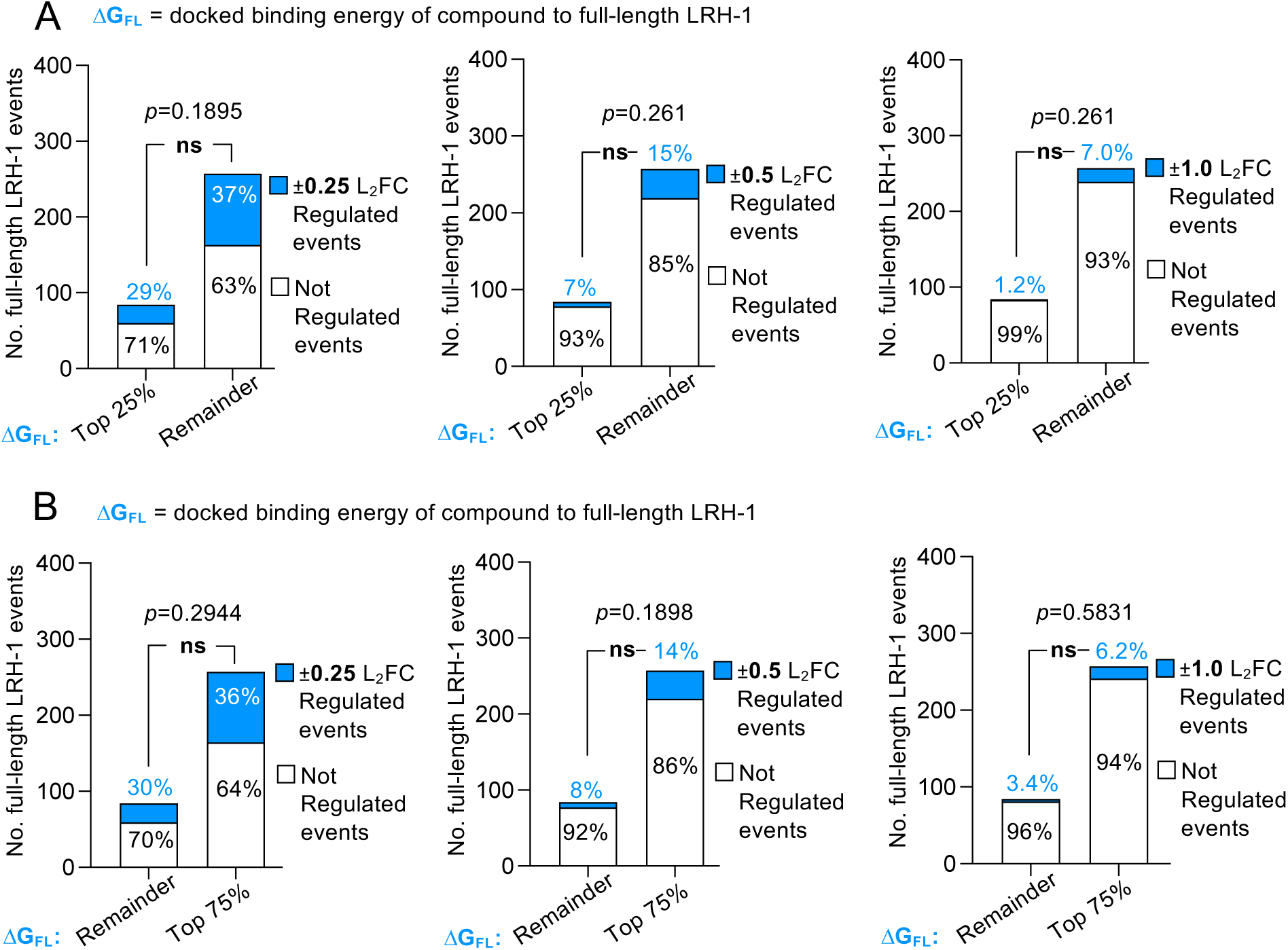
Computed compound binding energy to full-length LRH-1 does not in itself associate with regulation of full-length LRH-1 in cells. Contingency analyses of the frequency of LRH-1 regulation induced by all 57 compounds in all 341 full-length LRH-1 assayed events in cells (y-axis, each of the 341 wells in the high-throughput assay treated as an individual event, i.e. replicates not averaged) and regulation defined as meeting the indicated minimum log_2_ fold change cutoff (L_2_FC) compared to DMSO control, L_2_FC indicated in each panel. **Blue section** of bars is the number of regulated events (percentages indicate the percentage of regulated events), **white section** of bars is the number of unregulated wet lab assayed events (did not meet the minimum L_2_FC). **A**. Contingency analysis of the frequency of full-length LRH-1 regulation in cells, comparing compounds in the top quartile of the lowest docked binding energies (top 25% or the best 25% binding energies) to full-length LRH-1 vs. all other compounds, showing compounds with lower binding energies did not more frequently regulate LRH-1, at all L_2_FC cutoffs tested. **B.** Contingency analysis of the frequency of full-length LRH-1 regulation in cells, comparing compounds in the top 3 quartiles of lowest binding energy to full-length LRH-1 (Top 75% or the best 75% of binding energies) vs. all other compounds (Remainder), again showing compounds with lower binding energy to full-length LRH-1 did not more frequently regulate full length LRH-1 in cells (all statistics by Fisher’s exact test). *These data suggest the simple computed docked binding energy of compounds to full-length LRH-1 does not associate with the ability of a compound to regulate full length LRH-1 in cells*.

**Figure S5.**
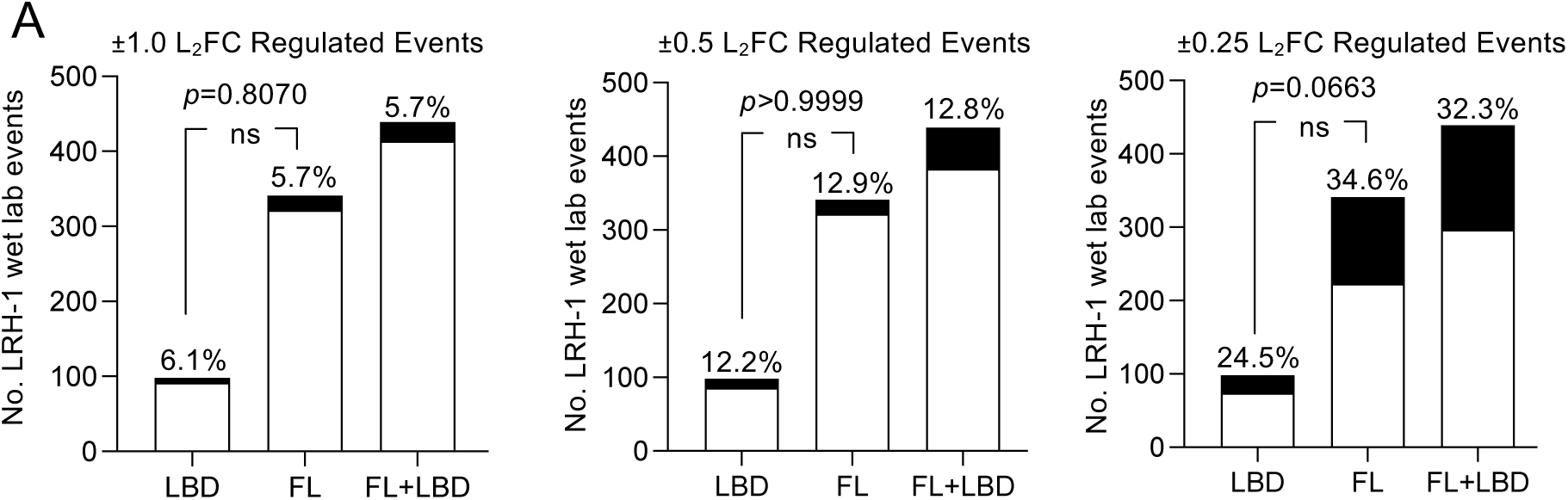
The 57 hit compounds had no inherent preference to regulate the isolated LBD vs. the full-length LRH-1 in cells when not separated into gorups. **A**. Frequency of indicated LRH-1 wet lab assays at ±1.0 L_2_FC cutoff (left), ±0.5 L_2_FC cutoff (middle) and ±0.25 L_2_FC cutoff (right). Contingency between the isolated LBD assay (LBD) *vs*. full-length LRH-1 assays in cells (FL) tested by Fisher’s exact, showing that without grouping the compounds, there is no inherent preference of these 57 compounds to regulate the isolated LBD or full-length LRH-1 assays. *These data suggest that without grouping the compounds into different classes, there is no difference in the frequency that compounds regulated the full-length vs. the isolated LBD assays*.

**Figure S7.**
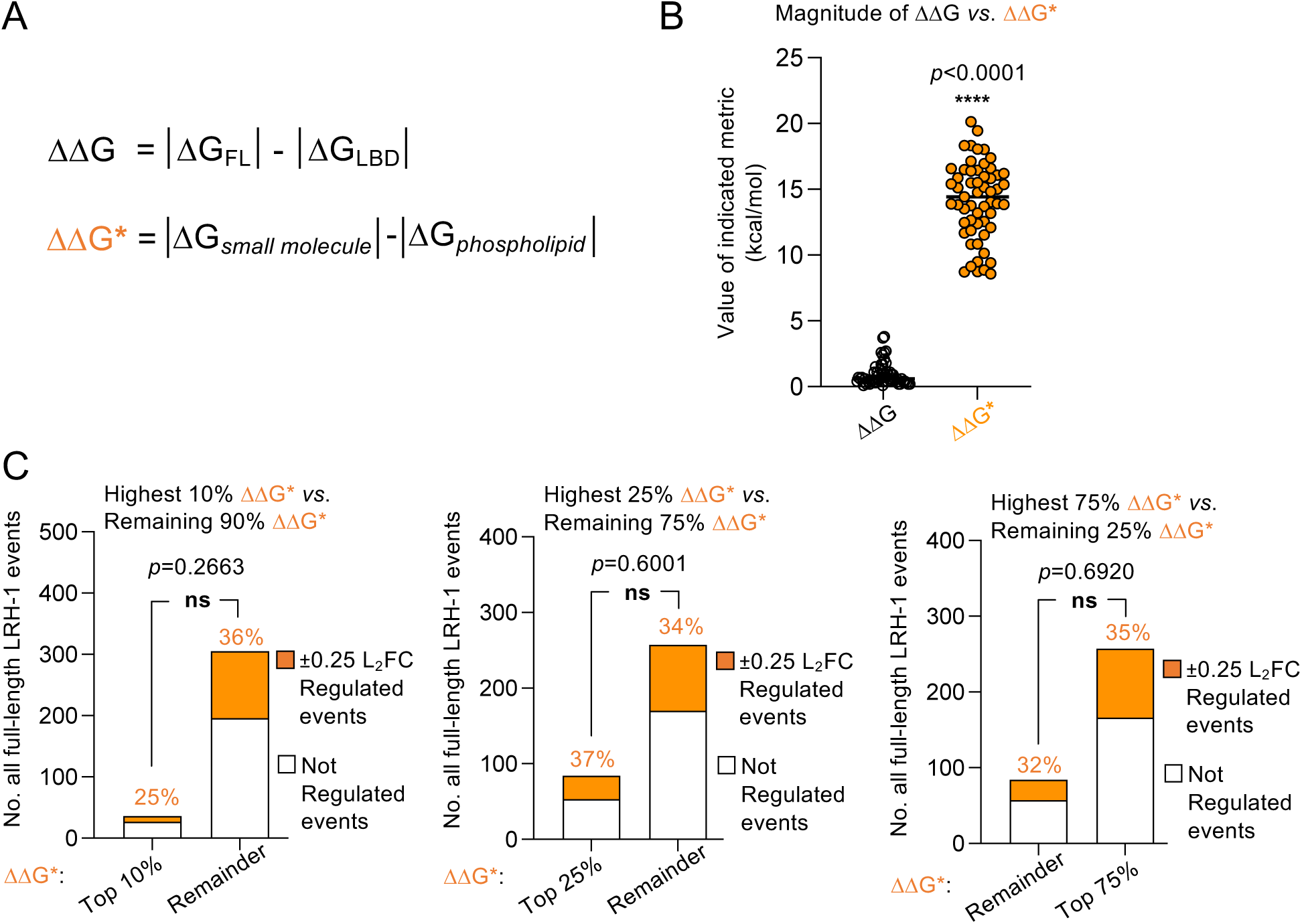
ΔΔG* does not associate with full-length LRH-1 activation in cells. **A**. To test if higher *ΔΔ*G values from small molecule-bound structures could explain the association of *ΔΔ*G with full-length LRH-1 activity in cells, we determined *ΔΔ*G* for all 57 compounds, *ΔΔ*G* was defined as the difference between the average ΔG for each compound bound to the 9 small-molecule co-crystal structures (*Δ*G*_small molecule_*) less the average ΔG bound to the 9 phospholipid co-crystal structures (*Δ*G*_phospholipid_*) for each of the 57 compounds **B.** Plotting the 57 values of *ΔΔ*G and 57 values of *ΔΔ*G* shows increased magnitude of *ΔΔ*G*, as expected. **C**. Contingency analyses showing the higher *ΔΔ*G* compounds did not more frequently regulate full-length LRH-1 in cells when compounds in the highest 10^th^ percentile (top 10%) *ΔΔ*G* were examined (*p*=0.2663), the highest quartile (top 25%) of *ΔΔ*G* compounds were examined (*p*=0.6001), or the highest 3 quartiles (Top 75%) of *ΔΔ*G* compounds (*p*=0.69203), by Fisher’s exact test. *These data suggest that ΔΔG* did not associate with the ability of these 57 compounds to regulate full-length LRH-1 in cell-based assays*.

